# A CRISPR/Cas9 screen reveals proteins at the endosome-Golgi interface that modulate cellular ASO activity

**DOI:** 10.1101/2024.12.17.628665

**Authors:** Liza Malong, Jessica Roskosch, Carolina Hager, Jean-Philippe Fortin, Roland Schmucki, Marinella G. Callow, Christian Weile, Valentina Romeo, Christoph Patsch, Scott Martin, Mike Costa, Zora Modrusan, Roberto Villaseñor, Erich Koller, Benjamin Haley, Anne Spang, Filip Roudnicky

## Abstract

Anti-sense oligonucleotides (ASOs) are modified synthetic single-stranded molecules with enhanced stability, activity, and bioavailability. They associate with RNA through sequence complementarity and can reduce or alter mRNA expression upon binding of splice site positions. To target RNA in the nucleus or cytoplasm, ASOs must cross membranes, a poorly understood process. We have performed an unbiased CRISPR/Cas9 knockout screen with a genetic splice reporter to identify genes that can increase or decrease ASOs activity, resulting in the most comprehensive catalog of ASOs-activity modifier genes. Distinct targets were uncovered, including AP1M1 and TBC1D23, linking ASOs activity to transport of cargo between the Golgi and endosomes. AP1M1 absence strongly increased ASO activity by delaying endosome-to-lysosome transport *in vitro* and *in vivo*. Prolonged ASOs residence time in the endosomal system may increase the likelihood of ASOs escape from this organelle before they reach lysosomes. This insight into AP1M1 role in ASOs trafficking suggests a way for enhancing the therapeutic efficacy of ASOs by manipulating the endolysosomal pathways.

## Introduction

Anti-sense oligonucleotides (ASOs) can modify the expression and/or splicing patterns of target RNAs, and have demonstrated effectiveness in the clinic, especially for undruggable targets^1^. Therapeutic ASOs are short single stranded oligonucleotides made from DNA or locked nucleic acid (LNA, with 2′-O-CH2-4′ linkage) with chemical modifications such as Phosphorothioate (PS) internucleosides^2^, or 2’-O-methoxyethyl (2’-MOE)^3^. ASOs can enter cells without assisted uptake in a process called gymnosis^4^, using different internalization pathways such as clathrin-mediated endocytosis, caveolin-mediated endocytosis, or dynamin-independent endocytosis^5^. Once inside the cell, ASOs reach early endosomes. Since ASOs do not contain any sorting signals, they remain in the endosome as it matures from early to late and recycles proteins to the plasma membrane and the Golgi apparatus^6^. Endosomes also receive cargo from the Golgi. Those proteins are either further transported to the membrane or, in the case of lysosomal hydrolases and resident proteins, routed to the lysosomes. Transport between the Golgi and endosomes is dependent on the clathrin adaptor complex 1 (AP-1)^7, 8, 9, 10^. The mature late endosome will then fuse with the lysosome to form an endolysosome in which the ASOs are degraded. In order to be effective, ASOs need to cross the endosomal membrane (“endosomal escape”) and enter the cytoplasm from which they then can also reach the nucleus^11^. However, endosomal escape is not very efficient; it is estimated that only 1-2% of ASO molecules are able to leave the endosome and most of the ASO content within the cell is ultimately trafficked to lysosomes for degradation^11, 12, 13, 14, 15, 16^. It is therefore of great importance to identify factors and pathways that, if modulated, would improve the escape of ASOs from endosomes into the cytoplasm and eventually into the nucleus, leading to more effective and, potentially, more durable activity profiles for this unique modality.

To reveal new modifiers of ASO trafficking to the cytoplasm and nucleus, we developed a surrogate genetic splice switch reporter system for which incorrect splicing of EGFP can be corrected with a splice-switch ASO (SS-ASO), serving as a functional, real-time readout of nuclear ASO activity^17^. We combined this reporter with unbiased whole genome CRISPR/Cas9 screening to identify, to our knowledge, the most comprehensive set of genes that increase or decrease the activity of the ASO. In particular, our CRISPR/Cas9 screen identified the cellular machinery involved in the transport between the Golgi and endosomes as being a rate-limiting step for ASO activity. The strongest negative modifier of ASOs function in our primary and validation screens was AP1M1, which is part of the clathrin adaptor AP1 complex. We further validated AP1M1 in distinct cell lines and an *in vivo* model, demonstrating that reduced expression of AP1M1 leads to degradation of TGN46 and a delay in the transport of DQ-BSA to lysosomes. We propose that targeted depletion of AP1M1 leads to a delay of ASO transport through the endosomal system, increased probability of ASO escape to the cytoplasm and, consequently, enhanced ASO activity.

## Results

### CRISPR/Cas9 KO whole genome pooled screen using the EGFP splice reporter identifies genes that modify ASO activity

We have developed an EGFP splice switching reporter to measure activity of splice switching ASOs by measuring directly and quantitatively (Fig. 1a, Extended data 1a-b) the correct splicing of EGFP. This system has been modified from a previously-published EGFP splicing reporter^18^ (EGFPsr), and shows no leakiness (Extended data 1a). Here, nuclear activity of a splice switch ASO (SS-ASO) results in exclusion of a β-globin intron-2 sequence placed within a mini-intron separating two exons used to create a full, in-frame copy of EGFP; explained in greater detail by ^17^ (Supplementary Table 1). To identify genes that modify cellular activity of ASOs, we subjected HEK293-Cas9-EGFPsr to treatment with 25 nM SS-ASO (Fig. 1b). Fluorescence-activated cell sorting (FACS) for the top 10% of GFP-positive (GFP+) and GFP-negative (GFP-) populations (Fig. 1b, Round1; Extended data 1c-d) was performed in two consecutive rounds allowing for one week of recovery after ASO treatment (Fig. 1b, round 2) to further enrich the positive and negative GFP cells (Fig. 1b). With this approach, we successfully identified sgRNAs (Supplementary Table 2) and the corresponding genes that, upon knockout, (Supplementary Table 3) increase (<u>SORT1 GFP+</u>, Fig. 1c; <u>SORT2 GFP+</u>, Extended data 2a) or decrease SS-ASO activity (<u>SORT1 GFP-</u>, Fig. 1d; <u>SORT2 GFP-</u>, Extended data 2b). Gene ontology enrichment analysis of the genes with fold change > 1.5 and p-adjusted-value < 0.05 for genes that increase the cellular activity of SS-ASO showed significant enrichment for molecular function of clathrin and cargo adaptor activity (<u>SORT1 GFP+</u>, Fig. 1e, upper diagram; Supplementary Table 4) and significant enrichment in cytoplasmic, Golgi, and clathrin/AP1 related proteins (<u>SORT1 GFP+</u>, Fig. 1e, lower diagram; Supplementary Table 4). Using the same cutoff for the genes that decreased the cellular activity of SS-ASO upon targeting, we found activity connected to histone acetyltransferase, iron and ascorbic acid and, interestingly, again clathrin and cargo adaptor activity (<u>SORT1 GFP-</u>, Fig. 1f, upper diagram, Supplementary Table 5). In addition, we observed significant enrichment in cellular components including endosomal transport, phosphatidylinositol-3-kinase complexes and lysosome (<u>SORT1 GFP-</u>, Fig. 1f, lower diagram, Supplementary Table 5). Of note, AP2M1, previously described to be involved in ASO function^5, 19^ was found in our screen. Together, our screen and analysis of both negative and positive modifiers of ASO activity found strong enrichment for vesicular trafficking, with particularly-high significance among clathrin-dependent adaptor complexes as well as the Golgi and endosomal systems.

**Fig. 1.**
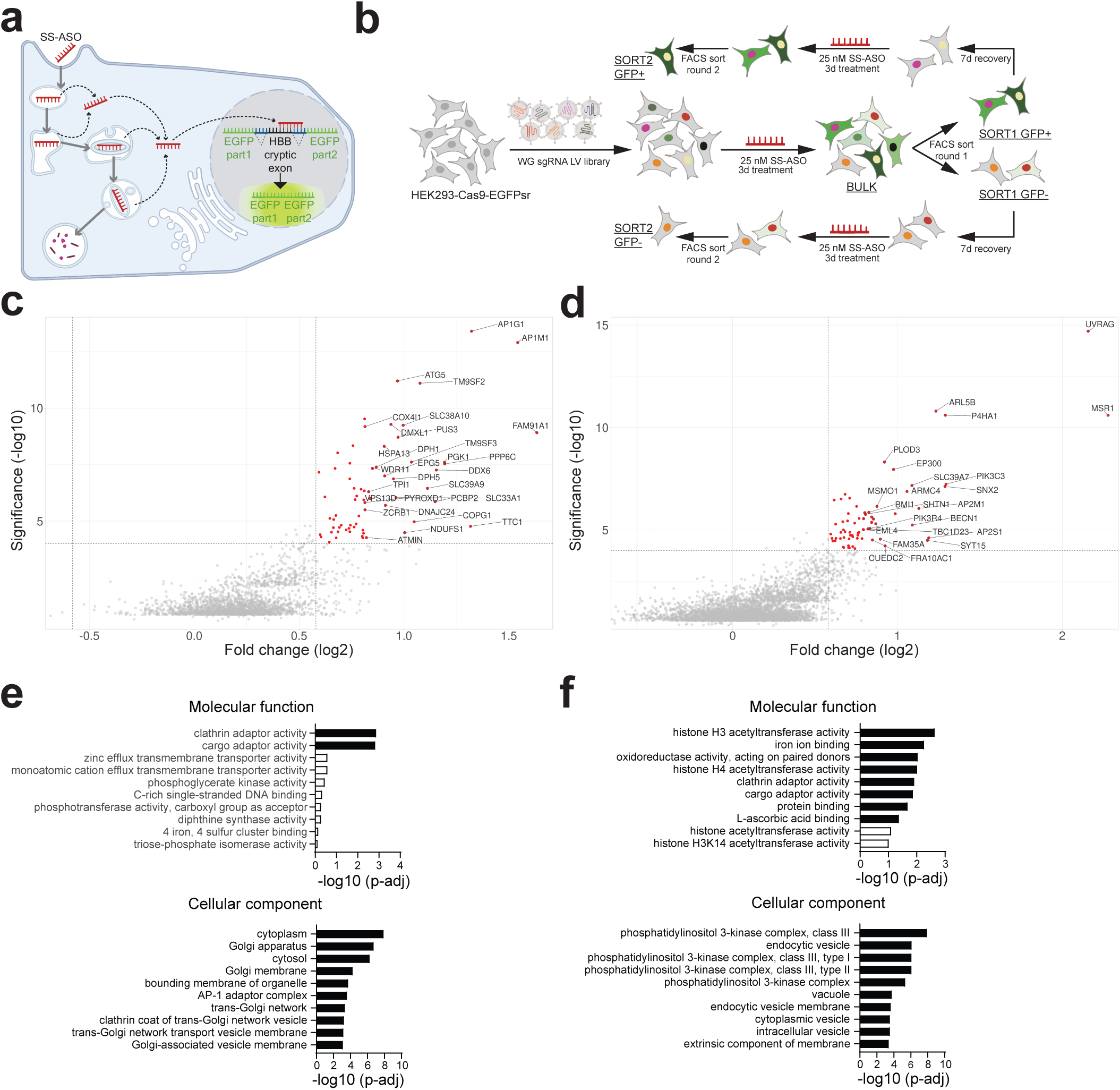
CRISPR/Cas9 KO whole genome pooled screen using the EGFP splice reporter identifies genes that modify ASO activity. **a**, Schematic representing splice switch reporter (HEK293-Cas9-EGFPsr) as a surrogate reporter for ASOs activity. ASO that enter the cell by gymnosis and escape the trafficking vesicles to the nucleus will bind to the cryptic exon of the splice reporter and cause the correct splicing and GFP expression. **b**, Schematic representing whole genome knockout CRISPR/Cas9 screen. HEK293-Cas9 cell line expressing HEK293-Cas9-EGFPsr were transduced with a whole genome sgRNA library, after selection cells were treated with SS-ASO (25nM), and top and bottom 10% GFP cells were FACS sorted (Round 1), after a week of recovery treatment was repeated followed by sorting using same conditions (Round 2). SgRNAs were deep-sequenced, and sequences were analyzed to identify sgRNAs that became enriched in sorted populations (<u>SORT1</u> and <u>SORT2</u>) relative to the unsorted population (<u>BULK</u>, populations that were NGS sequenced have an underline in the schematic). **c**, Volcano plot of genes from the comparison of Round1 of top 10% GFP+ vs unsorted population (<u>BULK</u>). **d**, Volcano plot of genes from the comparison of Round1 of top 10% GFP-vs unsorted population (<u>BULK</u>). Both Volcano plots show the significance -log10 (rho-value) versus the log2 (Fold change). Colored genes have a rho-value < 0.0001 and Fold change > 1.5. Top 30 genes sorted based on the fold change are labeled. **e**, Gene pathway analysis from significant genes (Fold change > 1.5 and p-adjusted value < 0.05) from the comparison of Round 1 of top 10% GFP+ (<u>SORT1 GFP+</u>) vs unsorted population (<u>BULK</u>) and **f,** from the comparison of Round1 of top 10% GFP (<u>SORT1 GFP-</u>) vs unsorted population (<u>BULK</u>) using g:Profiler with GO:MF (upper panel) and GO:CC (lower panel). Significant pathways are denoted with black bars (p-adjusted value < 0.05).

### Arrayed CRISPR screens validated ASO modifiers genes involved in retrograde trafficking and transport to and from Golgi compartment

We validated the primary hits from our knockout screen through lentiviral transduction of HEK293-Cas9-EGFPsr cells with multiple sgRNAs per gene (one sgRNA per well, 98 target genes in total) in an arrayed screening format (Fig. 2a-b, Supplementary Tables 6-7). Consistent with the effects in our primary, pooled screen, we were able to validate genes involved in vesicular trafficking such as AP1 clathrin adaptor complex (AP1M1, AP1G1), and coatomer genes (COPG1, COPB1, COPZ1). Using the same experimental workflow but this time with high content imaging as a readout for ASO activity, we observed that genes with the strongest effects were involved in trans-Golgi network (TGN) to endosomal system trafficking (Fig. 2c-d, Supplementary Table 6). Importantly, the AP1 adaptor complex, reported to be crucial for vesicular transport between endosomes and the TGN, was identified as one of the strongest hits in all three screening formats. We therefore focused our studies on the AP1 complex as a modifier of ASO trafficking. In addition, we were able to identify and validate genes that decreased ASO activity upon KO, including SNX2 and TBC1D23, which are believed to function in the same pathways as the AP1 and WDR11/Fam91a1 complexes^20, 21^ (Fig. 2e). Similar to WDR11/Fam91A, TBC1D23 has been implicated in the tethering of retrograde vesicles to the TGN^22^, yet they had opposing effects on ASO activity. We therefore decided to follow up on TBC1D23 as well.

**Fig. 2.**
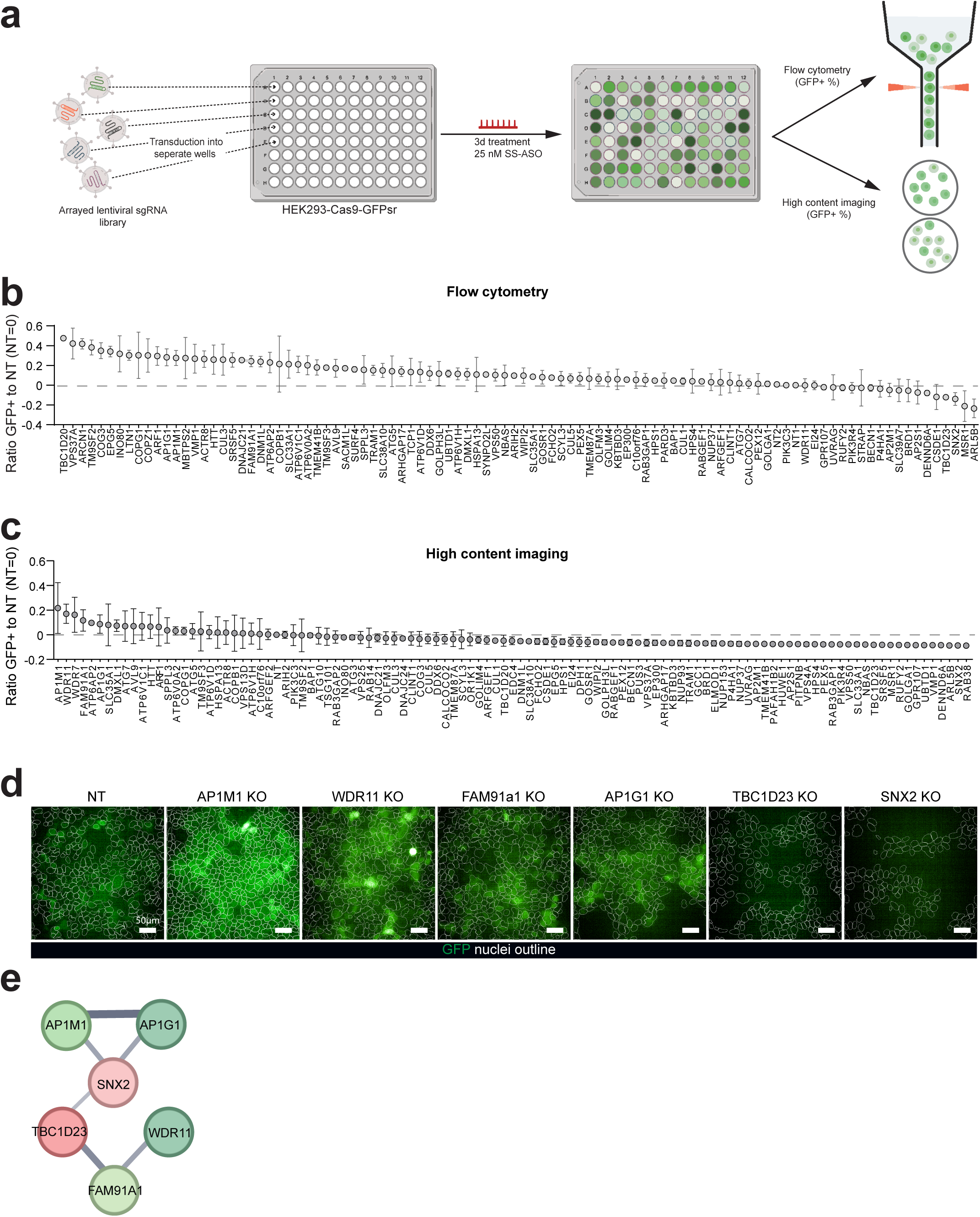
Arrayed screens validate genes involved in retrograde trafficking and transport to and from Golgi compartment. **a**, Schematic of arrayed validation screens. The sgRNA were delivered via lentiviral transduction. Following antibiotic selection, cells were treated with SS-ASO (25 nM, 3 days), and quantified the GFP+ expression by flow cytometry or high content imaging. **b**, Percentage of GFP+ cells was measured with flow cytometry. Screen was performed three times as independent experiments using 3 to 4 sgRNA per gene. Data is plotted as mean with SD and normalized to non-targeting sgRNA. **c**, Proportion of GFP+ cells was measured using high content imaging. Screen was performed as 2 independent experiments using 3 to 4 sgRNA per gene. Data is plotted as mean with SD. **d**, Representative images of the high content imaging results. Cellular segmentation was performed according to the Hoechst counterstaining and is depicted as white outlines. GFP is depicted in green. **e**, Protein-protein association network generated by STRING database performed on the hits presented on Fig. 2c. The disconnected nodes are not displayed. The thickness of the line between nodes represents the confidence of the interaction, with the thickest lines showing the highest confidence. The nodes are color-coded in red or green according to their effect on ASO activity. This analysis identifies the AP1 adapter complex, as well as the TBC1D123/Fam91a1/WDR1 complexes, outlining the importance of vesicular transport between endosome/golgi in determining ASO activity.

### AP1M1 and TBC1D23 are strong modifiers of ASO activity

We further investigated the roles of AP1M1 and TBC1D23 on ASO activity by generating monoclonal HEK293 knockouts (KOs) using CRISPR/Cas9 genome editing. Knockout efficiency was confirmed via next-generation sequencing (NGS) with CRISPResso2 analysis (Extended data 3a-b) and protein expression was assessed by Western blotting (Extended data 3c). We selected three AP1M1 KO and two TBC1D23 KO clones with successful allelic disruption evidenced by the absence of protein expression (Extended data 3a-c). To determine if the expected KO effects could be generalized beyond our splice switch context, we evaluated three distinct gapmers^3^ ASOs targeting CD81, MALAT1, or CERS2 in these AP1M1 and TBC1D23 KO cells. ASO efficacy was quantified by flow cytometry for CD81 and with qRT-PCR for MALAT1 and CERS2. Our results consistently showed enhanced ASO activity in AP1M1 KO cells and diminished activity in TBC1D23 KO lines, regardless of the ASO target or assay (Fig. 3a, Extended data 3d-e). To evaluate activity with a different cell line, we replicated the experiments in monoclonal U2OS cell lines with KOs for AP1M1 or TBC1D23, validated by NGS and Western blotting (Extended data 4a-c). Similar to HEK293 cell lines, AP1M1- and TBC1D23-KO U2OS cell lines showed significant positive and negative effects on ASO activity, respectively (Fig. 3b, Extended data 4d-e). Furthermore, time course experiments showed that the ASO activity occurs earlier in the AP1M1 KO lines compared to the parental populations (Fig. 3c-d). This was particularly evident in the U2OS context where target silencing was observed as early as 24 hours after ASO treatment in the AP1M1 KO *vs.* ∼48 hours in wild type cells (Fig. 3d). Combined, these results establish AP1M1 and TBC1D23 as significant modulators of ASO activity across different ASOs and cellular contexts.

**Fig. 3.**
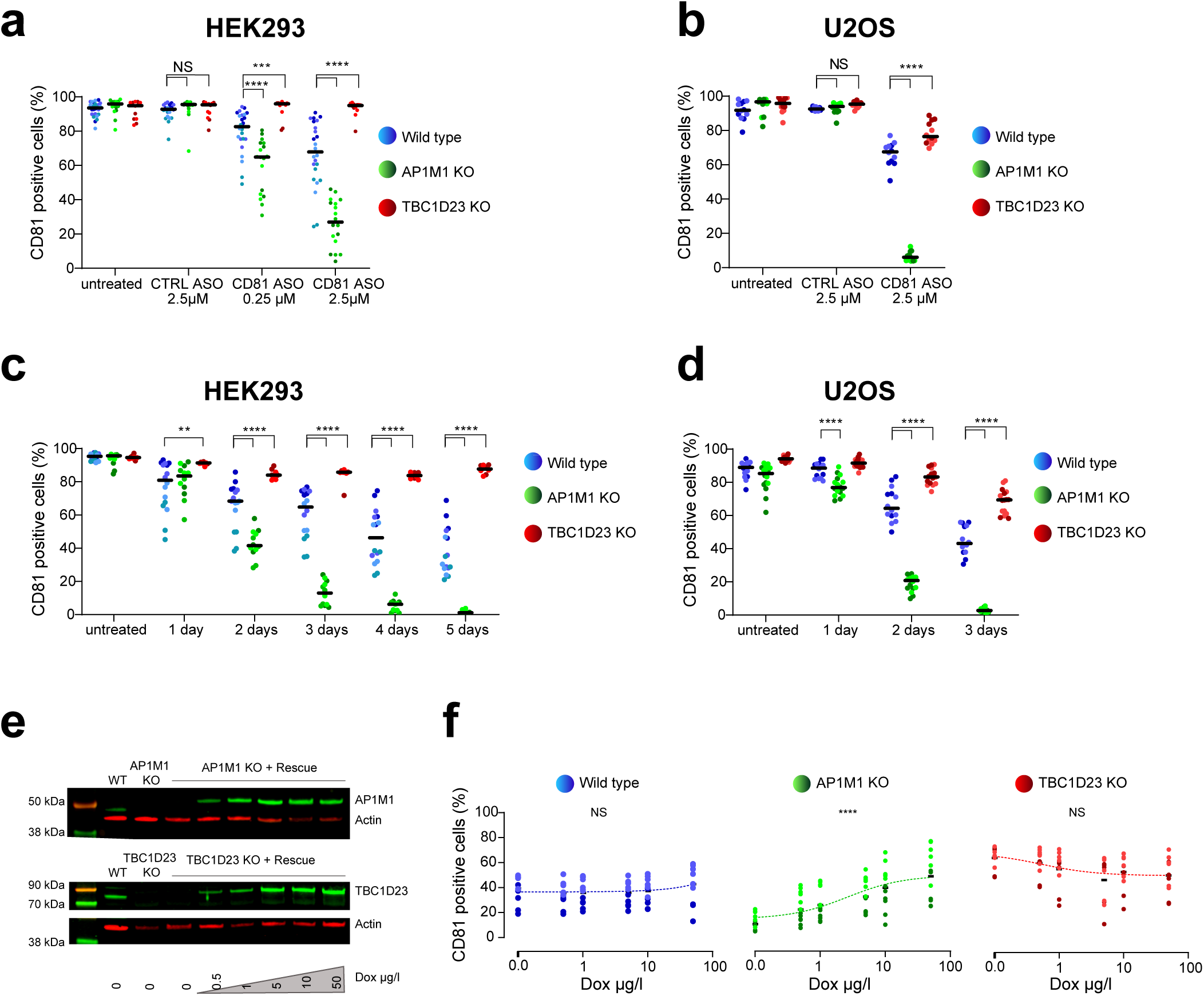
AP1M1 and TBC1D23 are strong modifiers of ASO activity. **a**, Percentage of CD81+ HEK293 were quantified by flow cytometry following 72h of treatment with an ASO against CD81 (0.25 or 2.5 µM), or a control non-targeting ASO (CTRL, 2.5 µM). **b**, Percentage of CD81+ U2OS were quantified by flow cytometry following 72h of treatment with an ASO against CD81 (0.25 and 2.5 µM), or a control non-targeting ASO (CTRL, 2.5 µM). **c**, Proportion of CD81+ HEK293 over time was quantified by flow cytometry, following 1 to 5 days time points of treatment with an ASO against CD81 (2.5 µM). **d**, Graph showing the proportion of CD81+ U2OS over time, as quantified by flow cytometry, following 1-3 days of treatment with an ASO against CD81 (2.5 µM). For the experiments above (**a-d**) horizontal lines show the median, each dot represents a technical replicate (n=2) from 2 or 3 independent experiments. Different color shades represent different clones. Statistical testing was performed with two-way ANOVA with Dunnett’s post-test (** p-value < 0.01; *** p-value < 0.001; **** p-value < 0.0001). **e**, Representative Western blot experiment of U2OS (AP1M1 or TBC1D23 KO) cells expressing the rescue construct (inducible expression of AP1M1 or TBC1D23), following doxycycline (Dox) treatment for 3 days. We used antibodies against AP1M1, TBC1D23 and ACTIN (loading control). The Western blot shows a dose-dependent re-expression of our genes of interest. **f**, Percentage of CD81+ in rescued U2OS as a function of Dox concentration, as quantified by flow cytometry, following 0.5 days of Dox treatment followed by 2.5 days of treatment with Dox and ASO against CD81 (2.5 µM). Dotted lines show the non-linear fitted curve (least squares regression method), each dot represents a technical replicate (n=2) from 3 independent experiments. Different shades of color represent different clones. Statistical testing was performed with a linear regression (R^2^ = 0.02, 0.31, 0.04 for wild type, AP1M1 KO and TBC1D23 KO, respectively; **** p-value < 0.0001).

### The effect of AP1M1 on ASO activity is specific and reversible

To ascertain the specificity and reversibility of our hits on ASO activity, we sought to rescue the phenotype observed in AP1M1 and TBC1D23 KO. For this, we used the Tet-On 3G system for doxycycline (Dox)-regulated expression of AP1M1 or TBC1D23 (Extended data 5a, Supplementary Table 8). U2OS cells with the corresponding gene KO were transfected with the rescue plasmids and stable integrants were generated by antibiotic selection. Dox titration demonstrated dose-dependent re-expression of AP1M1 and TBC1D23, as confirmed by Western blotting (Fig. 3e). Transgenic cells were pre-treated with Dox (half a day), followed by continued culture on Dox and ASOs (2.5 days). AP1M1 re-expression could abrogate ASO activity in a Dox concentration-dependent manner (CD81 ASO: Fig. 3f, Extended data 5b; CERS2 ASO: Extended data 5c-d). This was evidenced by a strong correlation between Dox levels and ASO efficacy for both CD81 and CERS2 ASO. TBC1D23 re-expression, while showing a trend, did not fully and significantly restore ASO activity to control levels in this context. Therefore, we decided to focus on the mechanisms by which AP1M1 modulates ASO activity.

### The effect of AP1M1 is downstream of gymnotic uptake

During the process of gymnosis, unformulated ASOs freely enter cells via endocytic vesicles^5^. ASO molecules then undergo endocytic trafficking, with most of them traveling to lysosomes (non-productive pathway), while only a small proportion escape the endosomal pathway and associate with their target in either the cytoplasm or the nucleus (productive pathway)^5, 19, 23^. Several surface receptors have been associated with gymnotic uptake of ASOs^5^, and we used whole-transcriptome RNA-seq to investigate whether perturbing AP1M1 results in differential expression of any known modifiers in this context. For this, we compared the two following pairs of cell lines: HEK293 cells wild type (WT) and AP1M1 KO (Supplementary Table 9) and U2OS wild type (WT) and AP1M1 KO (Supplementary Table 10). Differential expression (DE) analysis of the whole transcriptome RNA-seq identified few DE genes (|logFC| > 1, FDR < 0.05, logCPM > 1), and revealed there were no DE genes matching known gymnosis receptors (Extended data 6a, Supplementary Table 9 and 10). This shows that the phenotype observed in both cell lines upon AP1M1 KO is not driven by transcriptomic changes, and that no major rewiring of gymnotic uptake occurs in the absence of AP1M1. Beside gymnosis, another common way to deliver ASO is to use assisted uptake by packaging with cationic lipid nanoparticles (i.e. Lipofectamine RNAiMAX) prior to incubation with the cells. This route of administration leads to different uptake mechanisms and downstream delivery^5^. To determine whether alternative delivery methods were also dependent on AP1M1, we compared the impact of AP1M1 KO on ASOs delivered via gymnosis and lipid nanoparticle-mediated transfection. Notably, ASOs exhibit higher efficacy when delivered with transfection reagents such as Lipofectamine RNAiMAX (RNAiMAX), prompting us to conduct these experiments at an earlier time point (1 day for RNAiMAX versus 3 days for gymnosis). Our results indicated that in both HEK293 and U2OS cell lines, the activity of RNAiMAX-delivered ASOs was unaffected by AP1M1 KO (Fig. 4a-b, Extended data 6b-c), suggesting that AP1M1’s role is specific to the post-gymnotic uptake trafficking process.

**Fig. 4.**
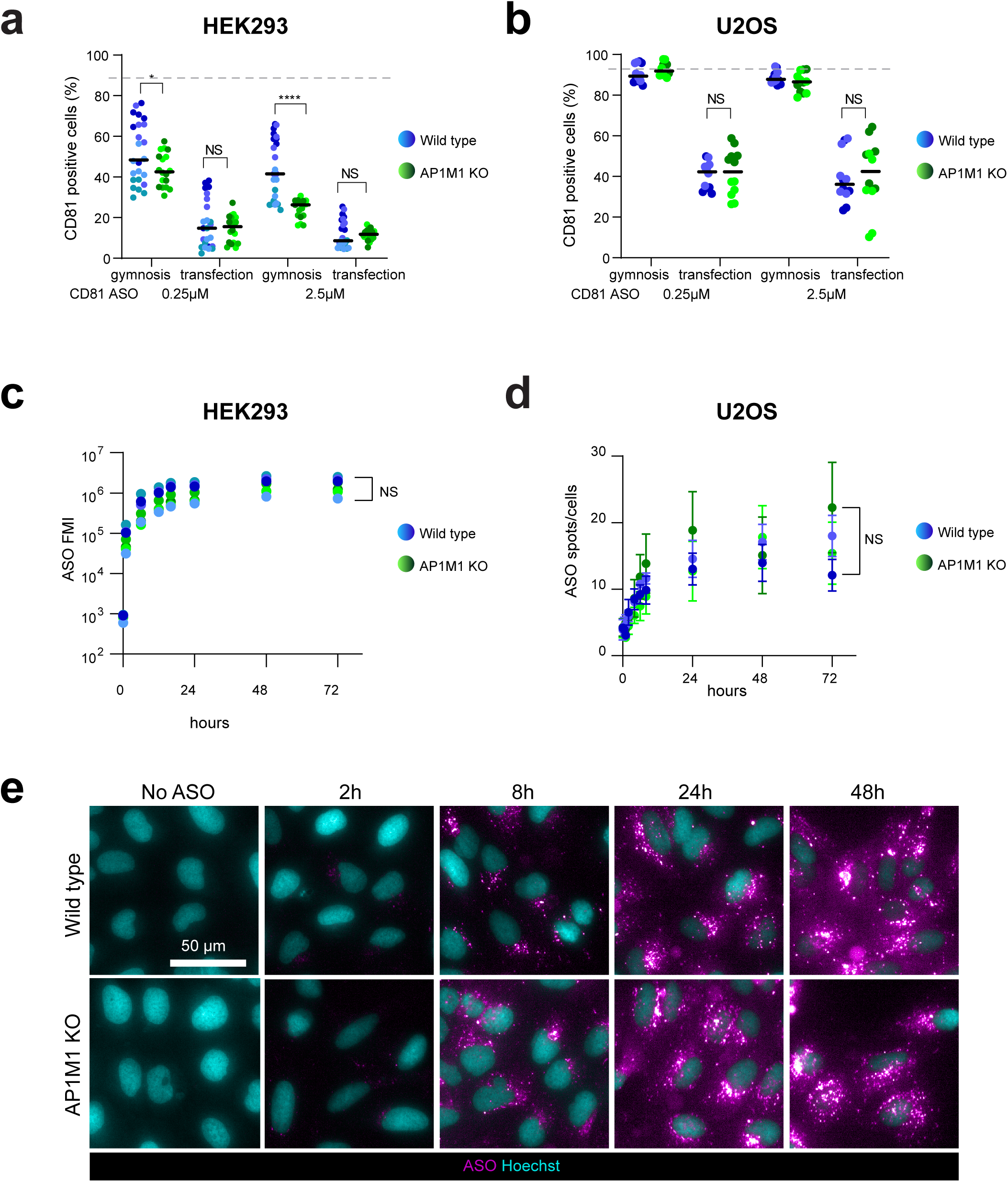
The effect of AP1M1 is downstream of gymnotic uptake. **a**, Percentage of CD81+ HEK293 cells was quantified by flow cytometry, following gymnotic or RNAiMAX delivery of CD81 ASO (0.25 and 2.5 µM) for 24 h. Horizontal lines show the median, each dot represents a technical replicate (n=2) from 3 independent experiments. Different color shades represent different clones. Statistical testing was performed with a two-way ANOVA with a Dunnett’s post-test (* p-value < 0.05; **** p-value < 0.0001). Dashed line depicts the percentage of CD81+ in untreated HEK293 clones. **b**, Percentage of CD81+ U2OS cells was quantified by flow cytometry, following gymnotic or RNAiMAX delivery of CD81 ASO (0.25 and 2.5 µM) for 24 h. Horizontal lines show the median, each dot represents a technical replicate (n=2) from 3 independent experiments. Different color shades represent different clones. Statistical testing was performed with a two-way ANOVA with a Dunnett’s post-test. Dashed line depicts the percentage of CD81+ in untreated U2OS clones. **c**, Fluorescence Mean Intensity (FMI, from Flow cytometry analysis) of HEK293 cells treated with a fluorescently labeled control non-targeting ASO (CTRL), over time (from 30 min to 72 h). Each dot represents the average value of a clone measured with 2 replicates from 2 independent experiments. Statistical analysis was performed with a two-way ANOVA with Dunnett’s post-test at 72 hours. **d**, Vesicles containing the fluorescently labeled ASO (spots) per U2OS cell were quantified from 30 min to 72 h by imaging. Each dot represents the average ± SEM of a clone measured with 2 replicates in 3 independent experiments. Statistical testing was performed with two-way ANOVA with Dunnett’s post-test at 72 hours. **e**, Representative images of the time course uptake experiment quantified in Fig. 4d. ASO is depicted with magenta while the nucleus is counterstained with hoechst and depicted in cyan. Scale bar corresponds to 50 µm.

To test whether AP1M1 loss affects ASO uptake, we measured ASO entry into cells using fluorescently-labeled ASOs. We confirmed that fluorescently labeled ASOs against CERS2 and MALAT1 maintain similar activity in cells compared to equivalent naked ASOs, validating the use of fluorescently labeled ASOs (Extended data 6d). Flow cytometry analysis revealed no significant difference in ASO uptake between AP1M1-KO HEK293 cell lines and control (WT) lines (Fig. 4c). In addition, these KO lines showed no difference in the uptake of other fluorescent markers, such as fluorescently labeled 40kDa dextran (Extended data 6e). Likewise, imaging analysis in U2OS lines also revealed no significant changes in ASO uptake in theAP1M1-KO context relative to wild-type (Fig. 4d-e). These findings suggest that AP1M1 KO does not influence the cellular pathways governing gymnotic ASO uptake. Instead, AP1M1 appears to modulate ASO activity through mechanisms related to intracellular trafficking and endosomal processing.

### Lysosomal function and morphology are undisrupted in AP1M1 KO cells

Following gymnotic uptake, most ASOs end up in lysosomes and undergo degradation^24^. Despite this being a major bottleneck of ASO therapies, ASO endosomal escape remains poorly understood^5, 14, 15, 25^. It is thought that a very small proportion of ASO molecules (< 1-2%) escape the endocytic pathway during their transit through the early or late endosomes^11, 15^. Given that the AP1 complex is known to facilitate the delivery and trafficking of lysosomal enzymes^26, 27^, we hypothesized that a disruption in AP1M1 might result in compromised lysosomal function, allowing ASOs to leak from lysosomes. We employed various live-cell lysosomal assays: Magic red (cathepsin B), Cytofix Red lysosomal stain (Cytofix Lyso), and SiR-lysosome stain (SiR Lyso), to assess lysosomal health but observed no significant differences between AP1M1 KO and wild type cell lines, apart from a slight increase in the number of Magic Red spots per cells (Fig. 5a-b).

**Fig. 5.**
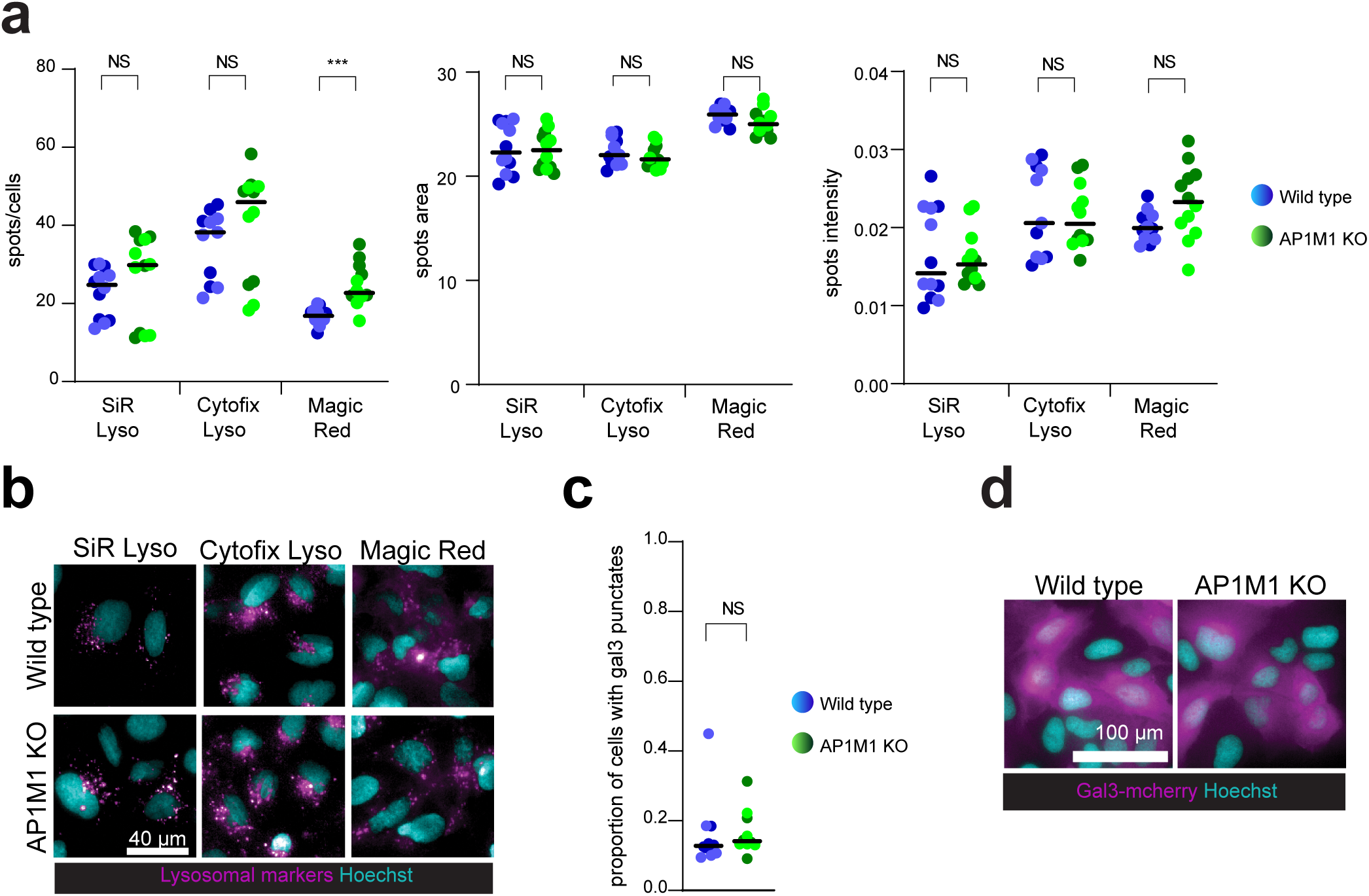
Lysosomal function and morphology is not disrupted in AP1M1 KO cells. **a**, The number of spots per cell, the area covered by these spots, and their intensity was quantified by imaging of U2OS cells treated with the following lysosomal markers: SiR Lysosome (SiR Lyso), Cytofix Red Lysosome (Cytofix Lyso) and Magic Red. Every condition had 2 technical replicates measured in 3 independent experiments. Different color shades represent different clones. Statistical testing was performed with the t-test (*** p-value < 0.001). **b**, Representative images of U2OS cells stained with lysosomal markers, as quantified in Fig. 5a. The different lysosomal markers are depicted in magenta, while the nucleus is counterstained with hoechst and depicted in cyan. Scale bar corresponds to 40 µm. **c,** Quantification of the proportion of cells that contain at least 1 galectin3-mCherry punctate (Gal3-mCherry) per cell was performed by imaging in U2OS clones overexpressing Gal3-mCherry. Horizontal lines show the median, 5 technical replicates per condition were measured in 2 independent experiments. Different color shades represent different clones. Statistical analysis was performed using an unpaired t-test. **d**, Representative images of Fig. 5c. Gal3-mCherry fluorescence is depicted in magenta, while the nucleus is counterstained with hoechst and depicted in cyan. Scale bar corresponds to 100 µm.

Moreover, we assessed lysosomal damage and permeability in AP1M1 KO in U2OS cell lines using galectin-3^28^ (Supplementary Table 8). Galectin-3 typically exhibits a diffuse cytoplasmic distribution but can relocalize to a punctate pattern as a result of recruitment to lysosomes upon increased permeability or lysosomal damage^29^. This relocalization can be induced by agents such as L-leucyl-L-leucine methyl ester (LLOMe), which we used as a positive control (Extended data 7a-b). AP1M1 KO cells did not display increased lysosomal permeability, as evidenced by the retention of a diffuse cytoplasmic distribution of galectin-3 regardless of AP1M1 expression (Fig. 5c-d; Extended data 7a-b). We conclude that AP1M1 KO cells have functional lysosomes, and suggested that the increased activity of ASOs may be due to an effect earlier in the trafficking pathway, specifically through more efficient endosomal escape.

### AP1M1 KO cells show perturbed trans-Golgi network

The AP1 complex is recognized as having a key role in transport between the trans-Golgi network and endosomes. More specifically, AP1M1 is involved in cargo sorting through the recognition of Yxx□ and [D/E]xxx[L/I] motifs^8, 30^, as well as acidic clusters^31^. Among those proteins are TGN46 (commonly used as a Trans-Golgi network (TGN) marker, involved in sorting of secretory proteins at the TGN^32^ and cation-dependent and cation-independent M6PR (receptor for lysosomal hydrolases, involved in lysosomal function and previously linked to ASO endosomal escape^33^). TGN46 localizes to the TGN as well as to endosomes. In AP1M1 KO cells, TGN46 levels appear to be generally lower compared to control cells, with the staining restricted to the TGN (Fig. 6a-c). This effect was specific for the AP1M1 KO and not due to a secondary effect because Dox-induced expression of AP1M1 restored TGN46 levels and distribution in AP1M1 KO cells (Fig. 6d-f).

**Fig. 6.**
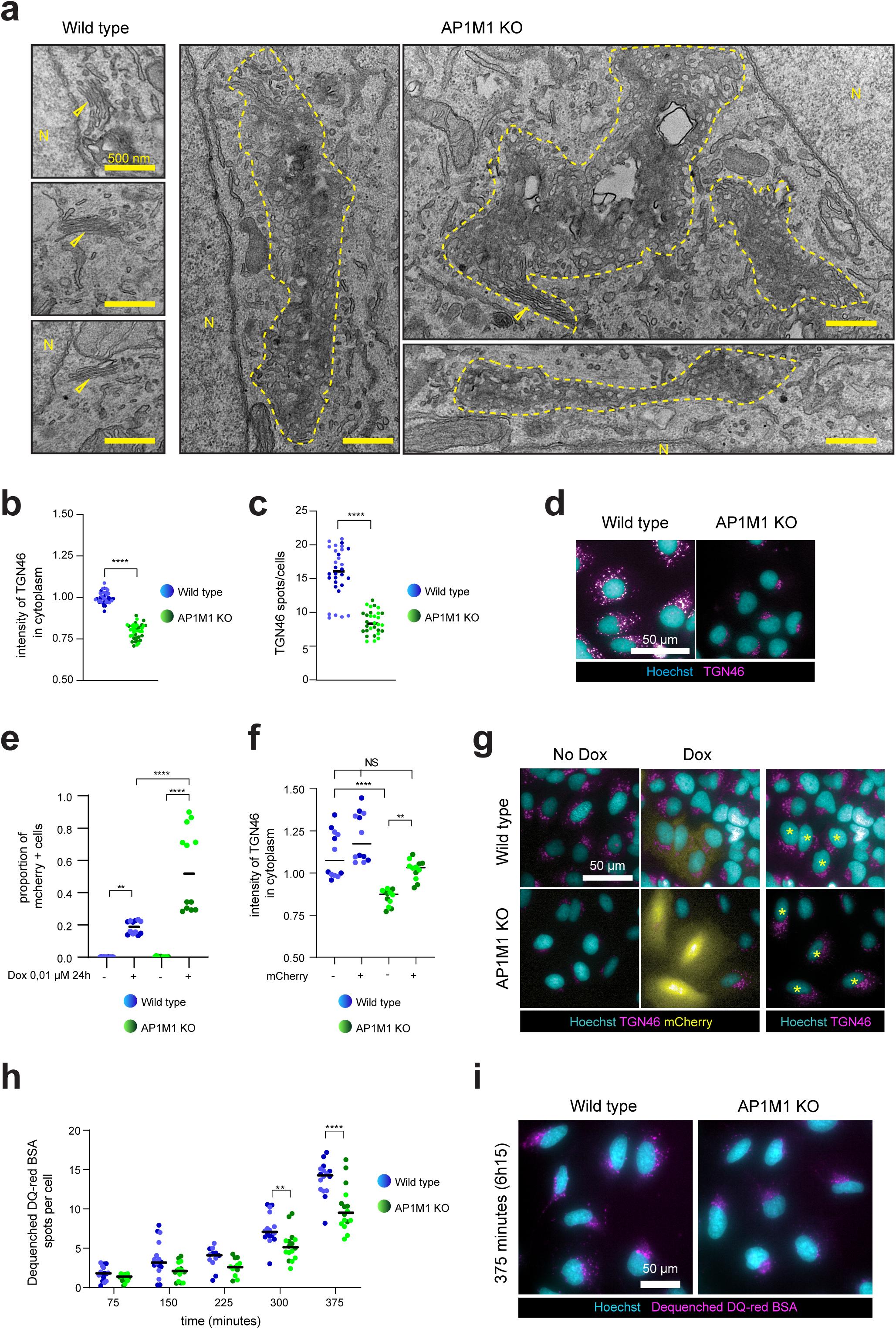
AP1M1 KO cells show perturbed trans-golgi network and show delayed transport of cargo to the lysosomes. **a**, Representative electron microscopy images of wild type and AP1M1 KO U2OS cells. Empty yellow arrowheads point to healthy golgi stacks. Dashed yellow lines surround Golgi anomalies. N depicts the nucleus. Golgi anomalies were rare in the wild type samples (n=11) but observed in most of the AP1M1 KO cells (n=30). Scale bar corresponds to 500 nm. **b-c**, TGN46 was immunostained in wild type and AP1M1 KO U2OS and quantified by imaging for staining intensity (**b**) and the number of spots per cell (**c**). Five technical replicates were measured in each of the 4 independent experiments. Different color shades represent different clones. Statistical testing was performed using an unpaired t-test (**** p-value < 0.0001). **d**, Representative images of TGN46 immunostaining (depicted in magenta), and nuclei counterstaining by Hoescht (depicted in cyan) in wild type and AP1M1 KO U2OS cells. **e,** Proportion of mCherry+ cells in wild-type and AP1M1 KO cells, expressing AP1M1 doxycycline-(Dox)inducible construct, were untreated or treated with Dox (0.01 µg/ml for 72 h) followed by quantification by imaging. In AP1M1 Dox-inducible construct, AP1M1 expression is linked to mCherry through an IRES element (Extended data 5a) and is regulated by Dox. Every condition had 2 technical replicates measured in 3 independent experiments. Different color shades represent different clones. Statistical testing was performed using one way ANOVA with Tukey post-test (** p-value < 0.01; **** p-value < 0.0001). **f**, Intensity of TGN46 staining was quantified by imaging in cytoplasm of wild type and AP1M1 KO cells expressing AP1M1 Dox-inducible construct, treated with Dox (0.01 µg/ml for 72 h). Every condition had 2 technical experiments measured in 3 independent experiments. Different color shades represent different clones. Statistical testing was performed using one way ANOVA with Tukey post-test (** p-value < 0.01; *** p-value < 0.001; **** p-value < 0.0001). **g**, Representative images of TGN46 immunostaining (depicted in magenta) in wild type and AP1M1 KO rescue cells treated with Dox (0.01 µg/ml for 72 h). MCherry (depicted in yellow) serves as a readout of the extent of the AP1M1 expression. Nuclei are counterstained with Hoechst (depicted in cyan). The far right panel shows the same images as the middle panel, without the mCherry signal, but with mCherry positive cells marked with a yellow asterisk. **h**, Number of de-quenched DQ-red BSA positive spots per cell (wild type and AP1M1 KO U2OS) was quantified by imaging over time (75 to 375 min) after treatment with DQ-red BSA. Horizontal line represents the median. Every condition had 2 technical replicates measured in 3 independent experiments. Different color shades represent different clones. Statistical testing was performed using one way ANOVA with Dunnett’s post-test (** p-value < 0.01; **** p-value < 0.0001). **i**, Representative images from Fig. 6h showing the fluorescent signal (dequenched DQ-red BSA in lysosomes, depicted in magenta) in wild type and AP1M1 KO cells. Nuclei are counterstained with Hoechst (depicted in cyan). Scale bar corresponds to 50 µm.

Our data suggests that Golgi morphology can be affected in AP1M1 KO cells. To this end, we performed electron microscopic analysis. Surprisingly, the Golgi morphology was not strongly altered (Fig. 6g), consistent with a recent report in which also only a delay in cargo export was reported^34^. We assume that other TGN export routes can partially overcome a large traffic jam in the Golgi.

### AP1M1 KO cells have delayed transport of cargo to the lysosomes

The reduction of TGN46 levels in AP1M1 KO cells could indicate that TGN46 can no longer be retrieved from endosomes to the TGN and therefore would be degraded in lysosomes, which we have shown to be functional. However, an increase in the protein load to be degraded by lysosomes may slow down endosome maturation and/or fusion of late endosomes with lysosomes. To investigate this possibility, we utilized DQ-red BSA, an albumin tracer conjugated to a quenched fluorophore that fluoresces upon lysosomal entry due to the pH-dependent activation of proteases which allows an unquenching mechanism (Extended data 8a). This approach ensures a high signal-to-noise ratio, as fluorescence is only emitted upon lysosomal arrival. Time-course experiments with this tracer revealed that AP1M1 KO cells have a significantly delayed endocytic cargo delivery to lysosomes, starting from 5 h post-incubation (Fig. 6 h-i). Consistent with our data, we did not find any difference in the general uptake of BSA, as seen by using Alexa Fluor 647 conjugated BSA (Extended data 8b), which is also in line with unaffected dextran and ASO uptake in AP1M1 KO cells (Extended data 6d). Our data indicate that endosome maturation or the fusion of late endosomes with lysosomes is delayed in AP1M1 KO cells. This delay would increase the residence time of ASOs in endosomes and thereby increase their chance to reach the cytoplasm and/or the nucleus.

### Ap1m1 expression modulates ASO activity *in vivo*

We further interrogated the role of AP1M1 in modifying ASO activity by testing its effect in an *in vivo* model of ASO activity. For these experiments, we used transgenic mKate2.ssEGFP.HBB mice^17^. These mice constitutively express a transgene containing mKate2 and the splice switching EGFP reporter interrupted by the hemoglobin intron (as used in our human *in vitro* CRISPR/Cas9 KO screen) under a CAG promoter in the Rosa26 safe harbor locus. Expression of the transgene was validated in all major organs of the mice, as well as the proportional and specific induction of splice switching events upon splice switch ASO treatment^17^. Since our *in vitro* results suggested that AP1M1 was particularly relevant for post-gymnotic entry, we first determined the conditions to produce optimal splice switching event/GFP expression following subcutaneous injection of the relevant naked ASO (gymnotic uptake). We subjected a cohort of adult homozygous transgenic animals to subcutaneous injection of splice switching or non-targeting control ASO (CTRL2, minimal effect on whole transcriptome as measured by total RNA-seq *in vitro*, Supplementary Table 11), and sacrificed animals after 3, 7 and 10 days. We measured a strong and consistent GFP signal in the liver and kidney at all time points using ELISA to quantitate GFP (Extended data 9a). In parallel, we performed experiments to assess the efficiency and liver-specificity of GalNAc conjugated siRNA against Ap1m1 or Ahsa1 (positive control for efficient knockdown, a previously validated liver target for GalNAc-siRNA^35^). We found that subcutaneous injection of a single dose of siRNA elicited a consistent and stable knockdown of both target genes 2 and 3 weeks post-injection, albeit with some variability in the case of Ap1m1 (Fig. 7b, Extended data 9b-d). Of note, we did not observe detrimental side effects of the treatment on the general physical state of the animals. We hence designed the experimental protocol as follows: at day 1, transgenic mice were injected (subcutaneously) with GalNAc-siRNA, at day 15 they were injected (subcutaneously) with ASOs, and at day 18 they were sacrificed (Fig. 7a). Knockdown of Ap1m1 and Ahsa1 mRNA, in liver (left panels) and kidney samples (right panels) collected at day 18, was quantified by qRT-PCR (Fig 7b).

**Fig. 7.**
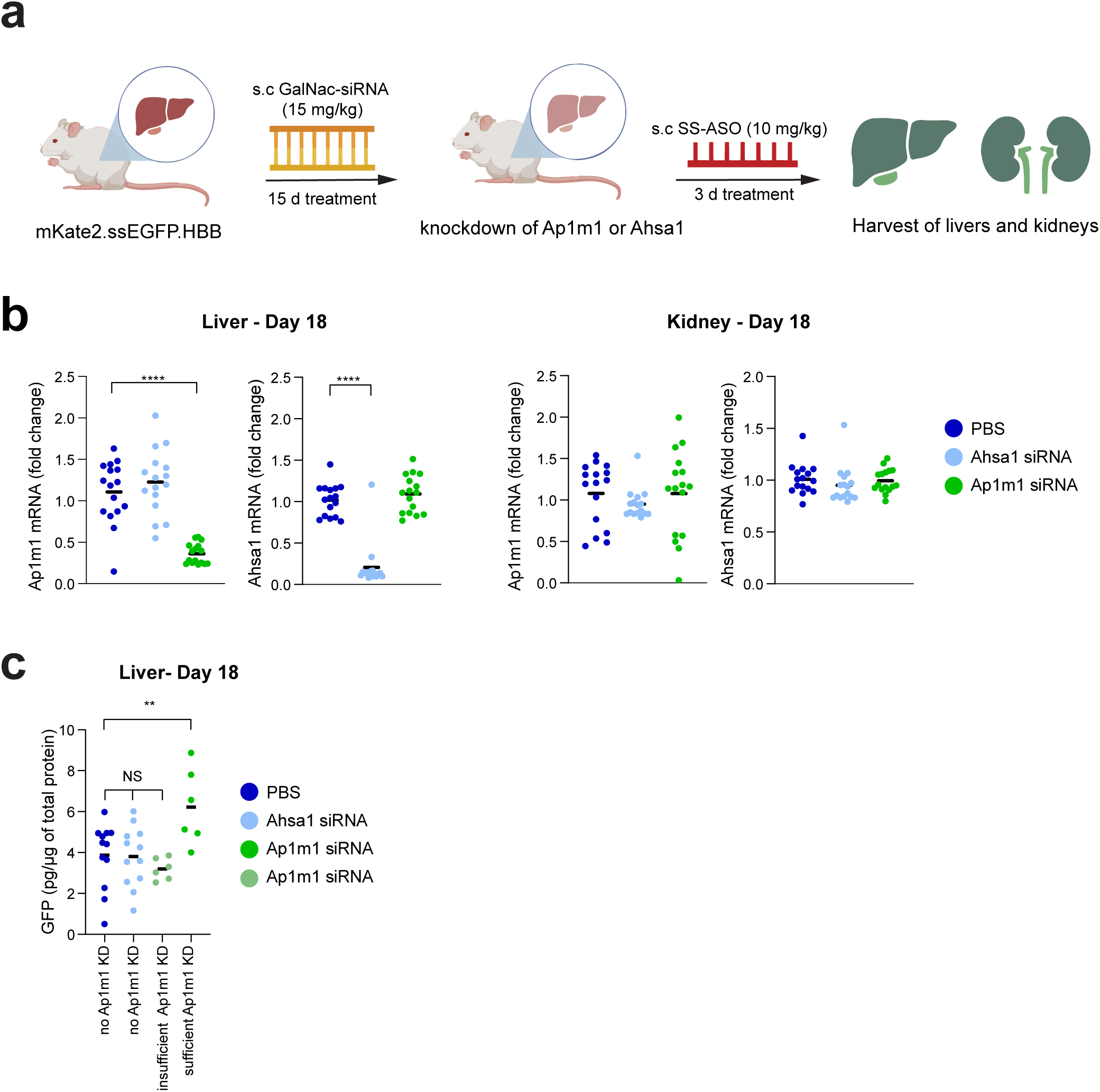
Ap1m1 expression modulates ASO activity *in vivo*. **a**, Schematic representing the *in-vivo* study design. Adult transgenic mKate2.ssEGFP.HBB mice carrying the splice reporter were injected subcutaneously with the GalNAc-siRNA targeting Ap1m1 or Ahsa1 (15 mg/kg) for 15 days before the administration of the SS-ASO, also via subcutaneous route. After 3 days, mice were sacrificed and the liver and kidneys were collected for subsequent analysis. **b,** Fold change of Ap1m1 and Ahsa1 mRNA, in liver (left panels) and kidney samples (right panels) collected at day 18, was quantified by qRT-PCR and normalized by housekeeping gene Gapdh. Horizontal lines show the mean, and each dot represents an animal. N= 16 animals per group, separated to 4 independent experiments. Statistical testing was performed with two-way ANOVA with Dunnett’s post-test (* p-value < 0.05; **** p-value < 0.0001). **c**, GFP protein expression, from the splice reporter (normalized by total protein level) was quantified at day 18. To account for the variability in Ap1m1 KD levels, the animals injected with Ap1m1 siRNA have been segregated into insufficient KD (less than one third knockdown) or sufficient KD (more than two third knockdown) groups. Each dot represents an animal, horizontal lines show the means. Analysis was performed with one-way ANOVA with Dunnett’s post-test (** p-value < 0.01).

Strikingly, we found that, similarly to our *in vitro* results, ASO activity significantly correlates with Ap1m1 levels in the liver of mice (Extended data 9e, Fig. 7c). In animals where Ap1m1 was sufficiently reduced (knockdown more than ⅔ of the transcripts), we found a significant increase in ASO activity as measured by GFP ELISA (Fig. 7c). This result, which recapitulates the *in vitro* screen findings showing a strong correlation between Ap1m1 expression and ASO activity, validates the role of Ap1m1 on ASO activity *in vivo*.

## Discussion

Activity of ASOs in a cellular system results from a combination of the intrinsic potency each ASO sequence (e.g. strength of binding to mRNA and recruitment of RNase H or displacement of splice factors) and the availability of ASO molecules to reach cytoplasmic and/or nuclear targets. The molecular mechanism by which ASOs escape the endosomal compartment is still elusive^25, 36, 37, 38, 39, 40, 41, 42, 43^. Indeed, studying ASO trafficking is challenging as conventional fluorescence microscopy can only identify the bulk of ASO molecules concentrated in otherwise non-productive, vesicular compartments, as opposed to the productive ones that are potentially dispersed as single molecules in the cytoplasmic or nuclear compartments^11, 12, 44^. To circumvent this confounding effect, we employed a functional read-out for ASO activity. Using this real-time based fluorescent reporter in conjunction with CRISPR/Cas9 genetic screening, yielded genes that increased or decreased ASO activity in a cellular context. Factors involved in membrane traffic, in particular within and between the Golgi and the endosomal system, were enriched within the GFP+ sorted fractions, where ASO activity was enhanced. These data indicate that escape from the endosomal system into the cytoplasm was likely to be a critical determinant of ASO activity. Subunits of the AP1 clathrin adaptor complex were among the strongest hits, and we subsequently validated the inhibitory effect AP1M1 has on ASO activity through orthogonal loss-of-function assays *in vitro* and *in vivo*. AP1 is known to be involved in transport of cargo between the TGN and endosomes^8, 10, 45^ , ^46^. Consistent with this notion, levels of the cargo adaptor TGN46 in the Golgi are reduced in AP1M1 KO cells. Similarly, in AP1 knockdown cells, late Golgi proteins leak into the endosomal system^45^. Moreover, reducing AP1 levels has been shown to slow recycling to the plasma membrane^47^. Thus, loss of AP1 appears to interfere with sorting in the endosomal system and, as a consequence, also slows endosomal maturation and transport to lysosomes^6^. Of note, decreased proteases activity in the lysosome is less likely to be the cause of the observed phenotype, as per the normal lysosomal protease markers in AP1M1 KO cells. We propose a model in which a delay in endosome maturation and fusion of late endosomes with lysosomes would provide ASOs with additional time to escape from endosomes, allowing them to reach the cytoplasm and, subsequently, the nucleus. This model is supported by our data showing that subunits of the V-ATPase and other proteins such as ATP6AP2, which are required for acidification and indispensable for lysosomal function, are likewise strong hits increasing ASO activity in our screens. Furthermore, a recent publication identified WDR91, a regulator of early to late endosomal maturation, as an ASO activity modulator^48^. Moreover, the small GTPase Arf1, which we discovered in our screen (increasing ASO activity), has been shown to play a role in endosomal transport as well as in transport through the Golgi^34,49,50,51^. In fact, the biosynthetic route through the Golgi is also important for ASO activity as highlighted by our identification of coatomer (COPI) components and other Golgi proteins as modifiers of ASO activity. Impairing transport through the Golgi will affect the downstream transport routes, such as the endolysosomal system. Altogether, our data strongly suggest that an efficient way to improve ASO activity is to modulate the trafficking pathway, in this case via AP1M1, before the ASO reaches the lysosomes. Interestingly, a similar effect to that of what we observed for ASOs in AP1M1 KO cells has been recently reported also for the transport of nanoparticles^52^. Nucleic acid therapies (mRNA vaccines, ASOs, siRNAs) are delivered either naked or with a delivery agent (LNP, nanoparticle) and they all rely on functional endosomes. This process is very inefficient, even when using delivery agents (∼2%)^12^. This opens the possibility that slowing down transport through the endosomal system could be a valid strategy to release not only ASOs but also other particles into the cytoplasm. Certain pathogens (e.g., *Mycobacterium*, *E. chaffeensis*, *N. risticii* or A. *phagocytophilum*) are able to slow down endosomal maturation and avoid fusion to lysosomes^53, 54^. Thus, a picture is emerging in which slowing down transport may promote endosomal escape. Modulating the speed of endosomal maturation may reduce infections (speeding it up) and increase the delivery of nucleic acid therapies (slowing it down).

In conclusion, our study highlights the critical role of the AP1 clathrin adaptor complex, particularly AP1M1, in modulating ASO activity. We clearly show that by disrupting the trafficking pathways, we can enhance the endosomal escape of ASOs, thereby increasing their therapeutic efficacy. Future research should focus on further elucidating the molecular mechanisms of ASO trafficking and exploring the therapeutic implications of modulating these pathways.

## Extended data legends

**Extended data 1. FACS sorting strategy for the CRISPR/Cas9 pooled screen.**

**a**, Percentage of GFP+ HEK293-Cas9-EGFPsr cells measured with flow cytometry after treatment with a SS-ASO or control ASO with 0, 40 nM or 5000 nM for 72h. **b**, Percentage of GFP+ HEK293-Cas9-EGFPsr cells measured with flow cytometry after treatment of SS-ASO (dose response: 0, 0.32, 1.6, 6, 40, 200, 1000 and 5000 nM) for 72h. **c**, Representative FACS sorting gates for cells (left panel), single cells (middle panel) and live cells (right panel). **d**, Representative example of a FACS sorting strategy for GFP- and GFP+ top 10% (upper panel) and the subsequent check of purity after sorting (lower panel). Arrows denote 10% of GFP- and GFP+ sorted populations.

**Extended data 2. CRISPR/Cas9 screens identify genes from intracellular trafficking to increase and decrease activity of ASOs. Enriched genes of Round2 FACS sorting.**

**a**, Volcano plot of genes from the comparison of Round2 of top 10% GFP+ vs unsorted population (Bulk). **b**, Volcano plot of genes from the comparison of Round2 of top 10% GFP-vs unsorted population (Bulk). Both Volcano plots show the significance -log10 (rho-value) versus the log2 (fold change). Colored genes have a significant rho-value lower than 0.0001 and fold change greater than 1.5. Top 30 genes sorted by fold change are labeled.

**Extended data 3. Generation and validation of monoclonal HEK293 KO cells.**

**a**, Percentage of genome modified NGS reads, quantified by CRISPResso2 analysis from AP1M1 KO and TBC1D23 KO HEK293 clonal lines. The editing frequency reflects the proportion of reads carrying indels in the sgRNA targeted sequence. **b**, Sequence of target sites of sgRNA (depicted in 5’-3’ orientation) for AP1M1 and TBC1D23 KO HEK293 clonal lines. The editing frequency is derived from NGS reads analyzed by CRISPResso2. The location of the sgRNA cut site is depicted by a dashed-line rectangle, while deletions are depicted in black and insertions are depicted in brown. The location of the sgRNA is indicated by a gray line beneath the sequence. **c**, Representative Western blot experiment of monoclonal HEK293 (AP1M1 or TBC1D23 KO) populations. We used antibodies against AP1M1, TBC1D23 and ACTIN (loading control). The Western blot confirms the loss of the proteins of interest upon knockout. **d**, CERS2 mRNA was quantified by qRT-PCR and normalized by house-keeping gene RPLP0, following 72h of treatment with an ASO against CERS2 or a control non-targeting ASO (CTRL, 1 µM each) in HEK293 clones. Data are presented as fold change relative to untreated. Horizontal lines show the median, every condition had 2 technical replicates measured in 3 independent experiments. Different color shades represent different clones. Statistical testing was performed using two-way ANOVA with Dunnett’s post-test (* p-value < 0.05; **** p-value < 0.0001). **e**, MALAT1 mRNA was quantified by qRT-PCR and normalized by house-keeping gene RPLP0, following 72h of treatment with an ASO against MALAT1 or a control non-targeting ASO (CTRL,1 µM each) in HEK293 clones. Data are presented as fold change relative to untreated. Horizontal lines show the median, every condition had 2 technical replicates measured in 3 independent experiments. Different color shades represent different clones. Statistical testing was performed using two-way ANOVA with Dunnett’s post-test (* p-value < 0.05).

**Extended data 4. Generation and validation of monoclonal U2OS KO cells.**

**a**, Percentage of genome modified NGS reads, quantified by CRISPResso2 analysis from AP1M1 KO and TBC1D23 KO U2OS clonal lines. The editing frequency reflects the proportion of reads carrying indels in the sgRNA targeted sequence. **b**, Sequence of target sites of sgRNA (depicted in 5’-3’ orientation) for AP1M1 and TBC1D23 KO U2OS clonal lines. The editing frequency is derived from NGS reads analyzed by CRISPResso2. The location of the sgRNA cut site is depicted by a dashed-line rectangle, while deletions are depicted in black and insertions are depicted in brown. The location of the sgRNA is indicated by a gray line beneath the sequence. **c**, Representative Western blot experiment of monoclonal U2OS (AP1M1 or TBC1D23 KO) populations. We used antibodies against AP1M1, TBC1D23 and ACTIN as a loading control. The Western blot confirms the loss of the proteins of interest upon knockout. **d**, CERS2 mRNA was quantified by qRT-PCR and normalized by house-keeping gene RPLP0, following 72h of treatment with an ASO against CERS2 (0.1 or 1 µM) or a control non-targeting ASO (CTRL, 1 µM) in HEK293 clones. Data are presented as fold change relative to untreated. Horizontal lines show the median, every condition had 2 technical replicates measured in 3 independent experiments. Different color shades represent different clones. Statistical testing was performed using Two-way ANOVA with Dunnett’s post-test (* p-value < 0.05; **** p-value < 0.0001). **e**, MALAT1 mRNA was quantified by qRT-PCR and normalized by house-keeping gene RPLP0, following 72h of treatment with an ASO against MALAT1 (0.1 or 1 µM) or a control non-targeting ASO (CTRL, 1 µM) in U2OS clonal lines. Data are presented as fold change relative to untreated. Horizontal lines show the median, every condition had 2 technical replicates measured in 3 independent experiments. Different color shades represent different clonal lines. Statistical testing was performed using Two-way ANOVA with Dunnett’s post-test (* p-value < 0.05; *** p-value < 0.001).

**Extended data 5. ASO activity upon reexpression of AP1M1 or TBC1D23.**

**a**. Schematic of the plasmid map containing the coding sequence for expression of AP1M1 or TBC1D23 in a doxycycline-(Dox) dependent manner. **b**, Percentage of CD81+ in rescued U2OS as a function of Dox treatment (0.05 µg/ml), was quantified by Flow cytometry, following 0.5 days of Dox treatment, followed by 2.5 days of treatment with Dox and control non-targeted ASO (CTRL, 1 µM). Each dot represents a technical replicate (n=2) from 3 independent experiments. Different shades of color represent different clones. Depicted are the control conditions relative to Fig. 3f. **c**, CERS2 mRNA was quantified by qRT-PCR and normalized by the expression of the housekeeping gene RPLP0 (depicted as fold change compared to untreated), in rescued U2OS as a function of Dox concentration, following 0.5 days of Dox treatment, followed by 2.5 days of treatment with Dox and with an ASO against CERS2 (1 µM). Dotted lines show the non-linear fitted curve (least squares regression method), every condition had 2 technical replicates measured in 3 independent experiments. Different color shades represent different clonal lines. Statistical testing was performed with a linear regression (R2 = 0.005, 0.46, 0.005 for wild type, AP1M1 KO and TBC1D23 KO, respectively; **** p-value < 0.0001). **d**, control conditions relative to Extended data 5c. Cells were treated with or without Dox treatment (0.05 µg/ml) and with a control non-targeting ASO (CTRL, 1 µM) instead of CERS2 ASO. Data are presented as fold change relative to untreated. Horizontal lines show the median, every condition had 2 technical replicates measured in 3 independent experiments. Different color shades represent different clonal lines. Statistical testing was performed using two-way ANOVA with Dunnett’s post-test.

**Extended data 6. Uptake of ASO by assisted uptake, activity of labeled ASOs and uptake of dextran.**

**a**, Volcano plots of log fold change expression between genes differentially expressed in HEK293 WT vs AP1M1 KO and U2OS WT vs AP1M1 KO. The genes with the highest log fold difference are labeled. **b**, Percentage of CD81+ HEK293 cells was quantified by flow cytometry, following gymnotic or RNAiMAX delivery of CD81 or control non-targeting ASO (CTRL, 0.25 and 2.5 µM) for 24 h. Horizontal lines show the median, each dot represents a technical replicate (n=2) from 3 independent experiments. Different shades of color represent different clonal lines. Statistical testing was performed with a two-way ANOVA with a Dunnett’s post-test (* p-value < 0.05; **** p-value < 0.0001). Graph includes the control conditions relative to Fig. 4a. **c**, Percentage of CD81+ U2OS cells, as quantified by flow cytometry, following gymnotic or RNAiMAX delivery of CD81 or control non-targeting ASO (CTRL, 0.25 and 2.5 µM) for 24 h. Horizontal lines show the median, each dot represents a technical replicate (n=2) from 3 independent experiments. Different shades of color represent different clones. Statistical testing was performed with a two-way ANOVA with a Dunnett’s post-test. Graph includes the control conditions relative to Fig. 4b. **d**, Relative mRNA expression, expressed as a percentage, was measured by qRT-PCR in HEK293 cells for both fluorescently conjugated and naked ASOs targeting MALAT1 and CERS2. Statistical testing was performed using one-way ANOVA. **e**, The number of vesicles containing the fluorescently labeled 40 kDa dextran per HEK293 cell was quantified by imaging and expressed as the mean, both with and without a 2-hour wortmannin pre-treatment. Horizontal lines show the median, each dot represents a technical replicate (n=2) from 3 independent experiments. Different shades of color represent different clonal lines. Statistical testing was performed using two-way ANOVA with Sidak’s post-test.

**Extended data 7. Lysosome leakiness upon LLOMe treatment and gene expression of WT and AP1M1 KO HEK293 and U2OS cells.**

**a**, Proportion of cells that contain at least 1 gal-3-mCherry punctate, following 3 h of treatment with 2 mM LLOMe was quantified by imaging. Horizontal lines show the median, and each dot represents a technical replicate (n=10) from two independent experiments. Different color shades represent different clonal lines. Statistical testing was performed with an unpaired t-test (**** p-value < 0.0001). **b**, Representative images of Extended data 7a. Gal3-mCherry fluorescence is depicted in magenta and nuclei are counterstained with Hoechst and depicted in cyan.

**Extended data 8. Schematic of entry and trafficking of DQ-red BSA and quantification of Alexa647-BSA entry.**

**a**, Schematic representing the uptake, endocytosis, trafficking, and subsequent de-quenching of DQ-red BSA upon arrival into the lysosomes. PH-dependent activation of proteases lead to de-quenching of DQ-red BSA **b**, AlexaFluor647-BSA positive spots per clones of AP1M1 and WT U2OS cells were quantified using imaging from 75 to 375 min after treatment with AlexaFluor647-BSA. Horizontal lines represent mean ± SEM. Every time point had 2 technical replicates measured in 3 independent experiments. Different color shades represent different clonal lines.

**Extended data 9. SiRNA mediated downregulation of Ahsa1 and Ap1m1 *in vivo.***

**a**, GFP protein levels, normalized by total protein amount, were measured by a GFP ELISA in livers and kidneys of animals sacrificed at day 3, 7, or 10. Each dot represents an animal. Horizontal line shows the median values per treatment from a single experiment (N=1 to 3 animals). **b**, mRNA fold change of Ap1m1 and Ahsa1 in liver samples collected at day 15, quantified by qRT-PCR and normalized by Gapdh (PBS treated group=1). Horizontal lines show the median, and each dot represents an animal. N= 2 to 3 animals per group from a single experiment. **c**, Quantification and representative image of Ap1m1 protein expression (red) from a Western Blot, normalized by albumin levels (green), in livers of mice sacrificed at day 15. Horizontal lines show the median, and each dot represents an animal. N = 2 to 3 animals per group from a single experiment. **d**, Quantification and representative image of Ap1m1 protein expression (red) from a Western Blot, normalized by albumin levels (green), in livers of mice collected at day 18. Lines show the mean, and each dot represents an animal. N= 16 animals per group, in 4 independent experiments. Statistical testing was performed with One-way ANOVA with Dunnett’s post-test (**** p-value < 0.01) **e**, Graph showing the levels of GFP protein, normalized by total protein amount, in livers of animals collected at day 18, plotted against the respective Ap1m1 mRNA level measured in the liver of each animal. Each dot represents an animal. n= 16 animals per group, in 4 independent experiments.

## sMethods

### Cell lines and culturing

The human Griptite HEK293-Cas9 line was obtained from Thermo Fisher. GNE293 line was obtained from Genentech Inc. cell repository. The knockout of AP1M1 and TBC1D23 in the U2OS line (HTB-96, Lot No. 58078676, ATCC) were obtained from Synthego Inc. HEK293 were cultured in DMEM supplemented with 1% Penicillin-Streptomycin (Thermo Fisher, Catalog No. 15140122), 10% FBS (VWR, 97068-085 or Gibco, Catalog No. A31605-02), 110 µg/ml Sodium Pyruvate (Sigma-Aldrich, Catalog No. S8636), Glutamax (Thermo Fisher, Catalog No. 35050061), 1x MEM Non-Essential Amino Acids (Thermo Fisher, Catalog No. 11140050). U2OS were cultured in Mccoy’s 5a medium (Gibco, Catalog No. 26600-023) complemented with 10% FBS (Gibco, Catalog No. A31605-02) and 1% Penicillin-Streptomycin (Thermo Fisher, Catalog No. 15140122). Cells were passaged using Accutase (StemCell Technologies, Catalog No. 07920) or TrypLE Express (Thermo Fisher, Catalog No. 12604013) or Trypsin-EDTA (0.05%, Thermo Fisher, Catalog No. 25300054) and re-plated as small clumps of cells at a dilution of 1:5 to 1:20. All cells were cultured in a humidified 37°C incubator set at 5% CO2 and passaged 1–3 times weekly.

### HEK293-Cas9 splice switch HBB EGFP reporter construction

Described in detail in^17^. Briefly, EGFP sequence is split at nucleotide (nc) 471 (counted from ATG-start-codon) with a β-globin intron-2 carrying a C to T mutation at nucleotide 1125 (counted from ATG-start codon; Supplementary Table 1). Intron sequence has been mutagenized at nc 1127-1128 from TA to GT (counted from ATG-start codon; intron site is at 657-658, Supplementary Table 1). Expression construct of split EGFP-HBB mutated intron has been synthesized and PCR-mutagenized from wild-type sequence (General Biosystems Inc.) and cloned in pcDNA3.1 (+) expression vector (Thermo Fisher, Catalog No. V79020). Vector was linearized with the restriction enzyme Nrul (Promega) and HEK293-Cas9 cells were nucleofected with 1 µg of plasmid using Amaxa SF Cell Line 4D Nucleofector kit (Lonza, Catalog No. V4XC-2032) with CW139 program. After 48 h post-nucleofection, the media was changed to DMEM with high glucose (Thermo Fisher, Catalog No. 11960-044) and sodium pyruvate (Sigma-Aldrich, Catalog No. S8636) and Glutamax (Thermo Fisher, Catalog No. 35050061) containing 600 µg/ml Hygromycin (Takara-Clontech, Catalog No. 631309) for selection. Selected Hygromycin-resistant population was sorted into single cells as follows: PBS -/- (Thermo Fisher, Catalog No. 14190094) washed cell suspension was filtered with 100 µm cell strainer (Corning, Catalog No. 352360). Cells were single-cell sorted with Cell Sorter SH800S (Sony Biotechnology) in poly-D-Lysine coated 96-well plates with DMEM medium supplemented with 20% FBS (Gibco, Catalog No. A31605-02), 5 µg/ml Blasticidin (Thermo Fisher, Catalog No. A1113803) and 600 µg/ml Hygromycin (Takara-Clontech, Catalog No. 631309).

### Pooled CRISPR/Cas9 sgRNA library design

The sgRNA whole genome library was complied from genome assembly GRCh37 (hg19) using a custom algorithm has been designed as in^55^.

### Pooled CRISPR sgRNA vector library production

The lentiviral library has been produced as in^56^ Briefly, 4 sub-libraries were produced (library1 sgRNAs: 18,991; library 2 sgRNAs: 18,990; library 3 sgRNA: 18,986; library 4 sgRNA: 18,975) as four arrayed oligo synthesis pools (Cellecta Inc.) and sub-cloned into a puromycin selectable lentivirus vector pLKO_SHC201 (Sigma-Aldrich) that contained an optimized sgRNA scaffold to generate pool library vectors.

### Lentivirus production for pooled and arrayed screens

Lentiviral particles for the pooled library have been produced as described in^57^. In brief, GNE293 line was plated in plates coated with 0.2% gelatine (Science Cell, Catalog No. 0423) and cultured in DMEM high glucose media (Thermo Fisher, Catalog No. 35050061) supplemented with MEM Non-Essential Amino Acids Solution (Thermo Fisher, Catalog No. 11140050) and Glutamax (Thermo Fisher, Catalog No. 35050061). Transfection mix comprised of sgRNA expression vectors of the pooled library, pCMVdR8.9 and pCMV-VSV-g (molar ratio 1:2.3:0.2) were transfected to GNE293 cells with the solution of Lipofectamine 2000 (Thermo Fisher, Catalog No.11668019) and Opti-MEM I reduced serum medium containing HEPES, NaHCO3, L-Glutamine (Thermo Fisher, Catalog No. 31985062). Six hours post transfection, the supernatant was removed and fresh media supplemented with DNase I 1 IU/ml (Qiagen, Catalog No. 79256), MgCl2 5 mM (Thermo Fisher, Catalog No. AM9530G) and 20 mM HEPES pH 7.2±0.1 (Thermo Fisher, Catalog No. 15630080) was added and kept overnight. The next day, media supplemented with 1% Penicillin-Streptomycin (Thermo Fisher, Catalog No. 15140122) was exchanged and after 24 h of culture, lentivirus containing supernatant was collected and filtered through a 45 µm pore filter (Nalgene Rapid Flow, Thermo Fisher, Catalog No. 166-0045). Filtered supernatant has been concentrated using Lenti-X solution (Clontech, Catalog No. 631231) and pelleted with 2100x g for 45 min at 4°C. Pellets have been resuspended with Hanks buffer (Ca^2+^, Mg^2+^; Thermo Fisher, 14025092) with 1% BSA low-endotoxin (Sigma-Aldrich, A2058) and frozen in dry ice/ethanol slush, and stored at -80°C. To produce lentiviruses containing individual sgRNAs in array format for all validation and follow-up experiments, synthesized sgRNA sequences were individually cloned into the BsmBI site of pCLIP-gRNA-EFS-Puro lentiviral plasmid (Transomic, Supplementary Table 6.). Plasmids were transformed into competent Stbl2™ E.coli (Thermo Fisher, Catalog No. 10268019); plated and individual colonies were selected and grown overnight. The identity of each sgRNA was confirmed by Sanger sequencing using a hU6-insert-F primer, 5’-ACTATCATATGCTTACCGTAAC-3’. The sgRNA library was packaged into lentiviral particles by co-transfecting HEK293T cells (Transomic) with the sgRNA plasmid, the pcDNA-delta8.91 packaging plasmid and a pCMV-VSV-G envelope plasmid. At 60 h post transfection, the supernatants were collected, aliquoted and frozen and -80°C. Viral titers were determined in HeLa cells by limiting dilutions under puromycin selection with colony counting. Sequences of sgRNA used for lentiviral transduction in arrays are listed in Supplementary Table 6 and 7.

### CRISPR/Cas9 lentiviral screen

In order to start the screen, each of the 4 lentiviral sub-library infectious titers were determined for HEK293-Cas9 cells as previously described^57^. Next, Forty-four million HEK293-Cas9 cells per biological replicate were seeded, and the next day were transduced with a whole genome human lentiviral library (4 sub-libraries mixed) at a multiplicity of infection (MOI) of 0.3. Three individual transductions were performed and handled separately in all consecutive steps. On day 3, a reference sample was collected, centrifuged (610x g, 10 min), supernatant removed and cell pellets snap frozen. On day 4, sgRNA expressing cells were selected by adding puromycin (1.5 µg/ml, Thermo Fisher, Catalog No. A1113803) throughout the study. On day 11 (replicate 1), 12 (replicate 2) and 13 (replicate 3) cells have been treated with 25 nM SS-ASO for 72 h. After 72 h, cells were dissociated from plates using Accutase (StemCell Technologies, Catalog No. AT104) and filtered through 30 µm filters to create a single cell suspension(Miltenyi Biotec, Catalog No. 130-098-458). Next, 38 million cells were centrifuged (610x g, 10 min), supernatant removed and cell pellets snap frozen; the rest of the cells were prepared in DMEM (phenol free, Thermo Fisher, Catalog No. 21063045) media for sorting. Sorting was performed in parallel on several BD FACSAria III equipped with a 70 µm size nozzle (BD Biosciences) using a 4-way purity precision mode. Cells were sorted into two bins based on GFP expression and lack of cell death staining (SYTOX™ Red Dead Cell Stain (Thermo Fisher, Catalog No. S34859): low (bottom ∼10%) and high (top ∼10%) with approximately 6 million cells per bin (from each high and low gates). Sorting was performed in cooled collection tubes with complete DMEM with 20% FBS and 25 mM HEPES. Ninety % of sorted cells were centrifuged (610x g, 10 min), supernatant removed and cell pellets snap frozen (Round1). Ten % of the sorted cells were reseeded and allowed to proliferate and recover from SS-ASO treatment to day 20 (replicate 1), day 21 (replicate 2), day 22 (replicate 3) and then retreated with 25 nM SS-ASO for 72 h. Exactly the same FACS sorting procedure as described above was repeated – cells were sorted into two bins based on GFP expression and lack of cell death staining (SYTOX™ Red Dead Cell Stain (Thermo Fisher, Catalog No. S34859): low (bottom ∼10%) and high (top ∼10%). Approximately 6 million cells per bin were sorted, cells were centrifuged (610x g, 10 min) supernatant removed and cell pellets snap frozen (Round2). FACS plots were generated using the Flowjo_V10 software (Flowjo).

### Genomic DNA preparation, NGS of CRISPR/Cas9 screen

Genomic DNA and NGS were performed as described in^57^. Briefly, genomic DNA was isolated from frozen pellets using Gentra Puregene Cell Kit (Qiagen, Catalog No. 158046) and dissolved in 100 µl low TE buffer (10 mM Tris-HCl + 0.1 mM EDTA). PCR amplification of 25 cycles with Phusion High-Fidelity PCR Master Mix (New England Biolabs, Catalog No. F532L) with PCR primers (forward: 5’-TCTTGTGGAAAGGACGAGGTACCG-3’; reverse: 5’-TCTACTATTCTTTCCCCTGCACTGT-3’) was performed. PCR purification was done following protocol with Agencourt AMPure XP beads on Biomek Fxp (both Beckman-Coulter, Catalog No. A63880). Next, 40 ng of PCR product was used in HyperPrep PCR kit (Roche, Catalog No. 07962363001) for end-repair, A-tailing and adapter ligation using xGenStubby adapters (IDT). The post-ligation purification was performed using Agencourt AMPure XP beads on Biomek Fxp (both Beckman-Coulter, Catalog No. A63880). Library was amplified using 2X KAPA HiFi HotStart ReadyMix (Roche, Catalog No. 09420398001) and compatible indexing primers (IDT) and purified using Agencourt AMPure XP beads on Biomek Fxp (both Beckman-Coulter, Catalog No. A63880). Samples were diluted and ran with a high output run on the Hiseq 2500 (Illumina). To generate sufficient number of reads, libraries were sequenced with 500-fold representation with the estimation of 70% sgRNA mapping frequency.

### Bioinformatic analysis of CRISPR/Cas9 screen

NGS read counts of the required depth were obtained and processed using the ScreenCounter alignment tool available in the CrisprVerse ecosystem^57, 58, 59^, a modification to previously published methods^60, 61^. Read-counts were normalized by using sgRNAs targeting a gold-standard set of non-essential genes^62^. More specifically, the distribution of counts was centered so that the median value for non-essential genes from^62^ is similar across samples. The normalized counts were transformed using log2(x+1) transformation. sgRNAs that had very low counts in the reference sample (less than 30 reads) were filtered out for subsequent analyses. For each sgRNA, a linear model on the log-normalized counts using limma-voom^63^ for each comparison of interest was calculated, generating a log-fold change between the 2 groups being compared, and a p-value associated with that guide. For each gene, the p-values of the 4 individual guides were combined into one summary p-value called the “rho statistic” using the RRAα framework implemented in the gCrisprTools R package^64^. The 4 log-fold changes of individual sgRNAs were summarized by taking the median log-fold change to get a representative log-fold change at the gene-level. SgRNA and gene fold-change on the 1st (day 11/12/13) and 2nd (day 20/21/22) FACS enrichment relative to the pre-sorted puromycin selected population. The complete results per sgRNA and per gene are presented in Supplementary Table 2 and 3, respectively.

### Gene set enrichment analysis of the CRISPR/Cas9 screen

g:Profiler^65^ was run using the web version (https://biit.cs.ut.ee/gprofiler/gost; accessed 1^st^ of August 2024). The annotation sources used were GO:BP, GO:MF, GO:CC, KEGG, REACTOME (REAC). Gene list for round 1 GFP+ vs bulk (Supplementary Table 4.) and GFP-vs bulk (Supplementary Table 5.) was obtained by application of Fold change > 1.5 and p-adjusted value < 0.05. Gene list was ordered based on fold change values. The run was based on the ordered query. Only annotated genes from Homo Sapiens were used for statistical testing. The control for multiple hypothesis testing was done with the g:profiler algorithm g:SCS with a significance threshold of 0.05.

### Individual Lentiviral sgRNA transduction for the arrayed screen and the generation of monoclonal KO populations

The human Griptite HEK293-Cas9 cells were seeded in 96 well plates in order to reach approx. 30% confluency on the day of viral transduction. On the day of lentivirus treatment, medium was replaced with fresh medium complemented with 8 µg/ml polybrene (Sigma-Aldrich, Catalog No. TR-1003-G). Lentivirus containing sgRNA (Transomic) was then added to the medium with MOI of 1 and incubated at 37℃ for 1 day. On the following day, puromycin (Thermo Fisher, Catalog No. A1113803) was added to the virus-containing medium to reach a final concentration of 1.1 µg/ml. To generate the monoclonal cell line populations, cells were selected for 2 weeks and we used the FACS sorter SH800S (Sony biotechnology) to generate single-cell colonies in 96 well plates. The clones were validated by NGS and CRISPRESSO2 analysis (see below), as well as Western blot analysis.

### Arrayed lentiviral screen and analysis

After 2 weeks of antibiotic selection, the transduced cells were treated with splice-switching ASO for 3 days (25 nM). For the FACS screen, cells were dissociated with TrypLE Express (Thermo Fisher, Catalog No. 12604013), centrifuged at 300x g for 5 min and then washed with PBS-/- (Thermo Fisher, Catalog No. 14190094). Specific binding was blocked by a 15 min incubation in the Fc block (1:50, BD Biosciences, Catalog No. 564219). Cells were incubated for 1 h with a CD81 antibody conjugated with APC (1:5, BD Pharmingen, Catalog No. 561958). Samples were processed with a MacsQuant Analyser 10 (Miltenyi Biotec). Data were analyzed using Flowjo_V10 software (Flowjo). For all genes that were represented by at least 3 sgRNAs (not causing cell death), the average of the 3 replicates was compiled. For the high content imaging screen, cells were fixed in Antigen Fix (Diapath) for 15 min at RT and incubated with Hoechst for 1 h at RT. GFP signal and Hoechst counterstaining were acquired with an Operetta CLS (Perkin Elmer). Data were analyzed with Harmony (Perkin Elmer). For all genes that were represented by at least 3 sgRNAs (not causing cell death, as seen by a minimum of 500 nuclei acquired in a field of 0.178 mm^2^), the average of the 2 replicates was calculated. A protein-protein interaction analysis was further performed using the String database (Version 12.0) with the 14 most significant genes from the high content imaging screen (Uncorrected Fischer p-value below 0.1). The network was computed with the following settings: full string network, with edges of the network representing the confidence of the interaction using all active interaction sources (minimum accepted confidence of interaction of 0.4).

### NGS sequencing and CRISPRESSO2 analysis

DNA extraction from cell pellets and subsequent NGS sequencing was performed by Microsynth AG Switzerland. DNA from eukaryotic cells was isolated with the Qiagen DNeasy Blood & Tissue 96 kit (Qiagen, Catalog No. 69504) following the manufacturer’s instructions. DNA was quantified by PreciseGreen (Lumiprobe, Catalog No. 12010). PCR primers (all 5’-3’ orientation) for TBC1D23 for HEK293 clonal lines (forward: ATTACCCAGTTGAGTGCATAGG, reverse: AATGCTGTGGAGGGTATAGCAG) for U2OS clonal lines (forward: TGGAAGCAGGAGGTTGTGAT, reverse: TCTAGATGCGGCATGCTCATAA) for AP1M1 for HEK293 clonal lines (forward: CAGGATCTTCCAGGCAGCAG, reverse: CTGTAGGGTGGCCAGGTATG) and for U2OS clonal lines (forward: CAGGATCTTCCAGGCAGCAG, reverse: CTGTAGGGTGGCCAGGTATG) + Illumina Tails was done using HOT FIREPol Multiplex Mix (Solis Biodyne, Catalog No. 04-36-00120) at an optimized annealing temperature and ideal PCR conditions (20 PCR cycles). First Step PCR products were purified using NGS Clean Beads (CleanNA, Catalog No. CNGS-0001). Second Step PCR was done using Nextera XT Index v2 Primers (Illumina) and HOT FIREPol Multiplex Mix (Solis Biodyne, Catalog No. 04-36-00120) doing 10 PCR cycles at optimized temperature. Second Step PCR products were purified using NGS Clean Beads (CleanNA, Catalog No. CNGS-0001), quantified using PreciseGreen (Lumiprobe, Catalog No. 12010) and equimolarly pooled. Optimized annealing temperatures were empirically determined by applying a temperature gradient and selecting the temperature that produced the most abundant yet specific product (single band). Subsequently, the PCR libraries were sequenced on an Illumina MiSeq platform using a v2 500 cycles kit. Sequencing data was analyzed using CRISPRESSO2^66,67^.

### RNA extractions for RNA-seq

Cultured cells (approx. 80% confluency) were washed with PBS-/- (Thermo Fisher, Catalog No. 14190094) and detached with TrypLE Express (Thermo Fisher, Catalog No. 12604013). Cells were centrifuged for 5 min at 300x g, medium was decanted, and cell pellets were frozen and stored at -80 ℃. When specified, cells were treated with non-targeting ASOs (1μM) for 3 days prior to sample preparation. RNA extraction from cell pellets, followed by library prep and RNA sequencing was performed by GENEWIZ Germany GmbH. Total RNA was extracted from frozen cell pellets using Qiagen RNeasy Mini kit following manufacturer’s instructions (Qiagen, Catalog No. 74106). RNA samples were quantified using Qubit 4.0 Fluorometer (Thermo Fisher) and RNA integrity was checked with RNA Kit on Agilent 5300 Fragment Analyzer (Agilent Technologies). RNA sequencing libraries were prepared using the NEBNext Ultra II RNA Library Prep Kit (New England Biolabs, Catalog No. E7775) for Illumina following manufacturer’s instructions. Briefly, mRNAs were first enriched with Oligo (dT) beads. Enriched mRNAs were fragmented for 15 min at 94°C. First strand and second strand cDNAs were subsequently synthesized, cDNA fragments were end repaired and adenylated at 3’ ends, and universal adapters were ligated to cDNA fragments, followed by index addition and library enrichment by limited-cycle PCR. Sequencing libraries were validated using the NGS fragment Kit on the Agilent 5300 Fragment Analyzer (both Agilent Technologies, Catalog No. DNF-473-0500), and quantified by using Qubit 4.0 Fluorometer (Thermo Fisher). The sequencing libraries were multiplexed and loaded on the Flow Cell of the NovaSeq 6000 instrument (Illumina) according to manufacturer’s instructions. The samples were sequenced using a 2×150 Pair-End (PE) configuration v1.5. Image analysis and base calling were conducted by the NovaSeq Control Software v1.7 on the NovaSeq 6000 instrument. Raw sequence data (.bcl files) generated from Illumina NovaSeq 6000 was converted into fastq files and de-multiplexed using Illumina bcl2fastq program version 2.20. One mismatch was allowed for index sequence identification.

### Whole transcriptome RNA-seq analysis

Base calling was performed with BCL to FASTQ file converter bcl2fastq2 version 2.20.0 (Illumina). FASTQ files were quality checked with the FastQC^68^ version https://depot.galaxyproject.org/singularity/fastqc:0.11.9--0. RNA-seq paired-end reads were mapped onto the human genome (build hg38) with read aligner STAR version https://depot.galaxyproject.org/singularity/star%3A2.7.10a--h9ee0642_0 using default mapping parameters^69^. Aligned reads were quality checked with MultiQC^70^ version https://depot.galaxyproject.org/singularity/multiqc%3A1.12--pyhdfd78af_0. Numbers of mapped reads for all RefSeq and/or Ensembl transcript variants of a gene were combined into a single value (i. e. count) assuming unstranded library by featureCounts^71^ version https://depot.galaxyproject.org/singularity/subread:2.0.1--hed695b0_0 and normalized as tpm (transcripts per million). For differential gene expression of the contrasts (1) HEK293 WT *versus* HEK293 AP1M1 KO, and (2) U2OS WT *versus* U2OS AP1M1 KO, we applied the edgeR^72^ algorithm and identified genes with absolute log2 fold change larger than 0.5 and Benjamini–Hochberg adjusted p value smaller than 0.05 as significantly changed.

### Piggybac overexpression for the generation of rescue cells (AP1M1 and TBC1D23) and Gal3-mCherry cells

U2OS cells were seeded to reach 40% confluency in 6 well plates in DMEM media containing supplements and maintained overnight. Transfection was performed using lipofectamin 3000 (Thermo Fisher, Catalog No. L3000001), following manufacturer’s instructions. Briefly, lipofectamine 3000 was diluted in OptiMEM reduced serum media (Gibco, Catalog No. 11058-021) following recommended proportions. A solution containing OptiMEM reduced serum media, P3000 enhancer (provided in the Lipofectamine 3000 kit), the Piggybac vector (Supplementary table 8) and Piggybac transposase vector (Hera BioLabs) vector at 3:1 ratio (total 1 µg DNA per well) was prepared and mixed with the Lipofectamine solution at a 1:1 ratio. The DNA:lipid complexes were added to the cells after a 5 minutes incubation at room temperature. The next day, the medium was replaced with fresh medium to wash away the Lipofectamine. Three days after transfection, stable cell lines were selected using 0.5 µg/ml Puromycin (Thermo Fisher, Catalog No. A1113803). Cells were maintained in puromycin-containing selection medium for a minimum of 2 weeks in order to achieve successful antibiotic selection.

### Flow cytometry analysis to assess CD81 ASO activity

Cells were dissociated with TrypLE Express (Thermo Fisher, Catalog No. 12604013), centrifuged at 300x g for 5 min and then washed with PBS-/- (Thermo Fisher, Catalog No. 14190094). Specific binding was blocked by a 15 min incubation in the Fc block (1:50, BD Biosciences, Catalog No. 564219). Cells were incubated for 1 h with a CD81 antibody conjugated with APC (1:5, BD Pharmingen, Catalog No. 561958), and 30 min with a fixable viability stain FVS450 (1:500, BD Biosciences, Catalog No. 562247). Finally, the cells were centrifuged and resuspended in 100 µL MACS buffer (Miltenyi Biotec, Catalog No. 130-091-221). FACS measurements were taken with a cytoFLEX LX device (Beckman Coulter).

### ASO transfection

ASO treatment with Lipofectamine RNAiMAX (Thermo Fisher, Catalog No. 13778075) was performed according to the manufacturer’s instructions. Briefly, ASO/RNAiMAX solution was prepared by incubating a solution of ASO diluted in OptiMEM (Gibco, Catalog No. 11058-021) with a solution of RNAiMAX diluted in OptiMEM for 5 minutes at room temperature. We followed the manufacturer’s ratio suggestion according to the plate type used. ASO/lipid complexes were then incubated on approx. 60% confluent cells for 1 day before the cells were processed for ASO activity readout.

### RNA extraction, reverse transcription, and qRT-PCR for cell lines to assess CERS2 and MALAT1 ASO activity

Following ASO treatment, RNA extraction was performed with the RNeasy kit (Qiagen, Catalog No. 74181) following manufacturer’s instructions. Briefly, cells from 96 wells plates were lysed with 80 µl of RLT lysis buffer provided in the kit, complemented with DNAse I (Qiagen, Catalog No. 79256), and one volume of ethanol 70% was added to the lysate. This solution was loaded into the RNA extraction column and extraction was subsequently performed following manufacturer’s instructions. Reverse transcription was performed using components of the Cells-to-CT Bulk RT Reagents (Thermo Fisher, Catalog No. 4391852C) and following manufacturer’s instructions. Briefly, each reaction contained approx. 10 ng of RNA freshly extracted, RT buffer (1X), and RT enzyme mix (1X), in a final volume of RNase free water (Thermo Fisher, Catalog No. AM9937) of 50 µl. The thermocycler program was the following: 10 minutes at 42℃, 1 hour at 37℃, hold at 4℃. The expression of mRNA was quantified by TaqMan qRT-PCR with the QuantStudio 7 Flex real-time PCR system and the 2-DDCt method (Livak et al 2001). TaqMan probe/primer sets for human MALAT1 (Thermo Fisher, Catalog No. Hs00273907_s1) and human CERS2 (Thermo Fisher, Catalog No. Hs00371958_g1r). Each reaction was multiplexed with human reference gene hRPLPO (Thermo Fisher, Catalog No. 4310853). Expressions were normalized to the expression of the reference gene. The thermocycler Quantstudio Flex 12k Flex (Thermo Fisher) was used using a 40 cycles program.

### Immunofluorescence

Cells were fixed before reaching 100% confluency by a 15 min incubation in Antigen Fix (Diapath, Catalog No. P0016). Fixed cells were permeabilized in PBS-Triton X-100 0.1% for 15 min and blocked in BSA 2% for 1 h. Cells were then incubated in the following primary antibody against TGN46 (Bio-Rad Laboratories, Catalog No. AHP500g) overnight at 4℃. The next day, cells were washed in PBS-/- and incubated in corresponding secondary antibodies, as well as Hoechst and, when needed, phalloidin-alexa fluor 488 for 1 h at RT. Cells were further washed and acquisition was performed with an Operetta CLS (Perkin Elmer) in PBS-/-.

### High-content imaging

High content imaging was performed in imaging plates (Corning, Catalog No. 3904). All high content imaging was performed with an Operetta CLS (Perkin Elmer) equipped with an LED light source and analyzed with the built-in image analysis software, Harmony (Perkin Elmer). Images were acquired with a 40X or 63X lens. The equipment is fitted with the following filter sets (430-500/ 500-550/ 570-650/ 655-760).

### Transmission electron microscopy

For electron microscopy, cells were grown on glass coverslips. Cells were fixed by adding an equal volume of a warm solution containing 4% PFA (Electron Microscopy Sciences, Catalog No. 15710) and 5% glutaraldehyde (GA, Electron Microscopy Sciences, Catalog No. 16310) in 0.1M PIPES buffer (Sigma, Catalog No. P6757) with 2 mM CaCl2 (final concentration in the well was 2% PGA and 2.5% GA). After 15 min of fixation at RT, the solution was replaced by a fresh solution containing 2% PFA and 2.5% GA in 0.1 M PIPES buffer with 2 mM CaCl2 (pH 7-7.3). Cells were incubated in that solution for 2 h at room temperature, then 16 h at 4℃. After the fixation, cells were washed 3 times for 10 min with a cold 0.1 M PIPES buffer with 2 mM CaCl2, and delivered to the BioEM facility of University of Basel to undergo embedding and imaging as follows. Briefly, fixed cells were rinsed in Cacodylate buffer (0.1 M, pH 7.3, Sigma-Aldrich, Catalog No. 233854, further referred to as CaCo) three times and further fixed in a solution of CaCo containing 1% osmium tetroxide (Electron Microscopy Sciences, Catalog No. 19100), 0.8% potassium ferrocyanide for 1 h at 4℃. Following further washes in CaCo and water, cells were stained in a CaCo solution containing 1% uranyl acetate (Electron Microscopy Sciences, Catalog No. 22400) for 1h at 4℃ in the dark. Cells were further washed in water and dehydrated by incubation in ethanol solutions of increasing concentration, up to reaching absolute ethanol (30%, 50%, 75%, 96%, 100%, 100%, 100%, 100%). Dehydrated cells were washed in acetone (Electron Microscopy Sciences, Catalog No. 15056), incubated in resin/acetone, and then in Epon 812 resin (Electron Microscopy Sciences, Catalog No. 14120) overnight. Resin polymerization was achieved at 60℃ for 48 h on BEEM capsules (Electron Microscopy Sciences, Catalog No. 70010-B) filled with Epon. Following the polymerization, coverslips were removed from the Epon block and cells were transferred from the coverslip to the block surface. A diamond blade was used to cut 70 nm thin sections, which were collected on formvar-carbon coated copper grids. Grids were further stained with uranyl acetate and Reynolds’s lead citrate (prepared by following protoco from Reynolds 1963l). Images were acquired using a FEI Technai G2 Spirit transmission electron microscope and a EMSIS Veleta camera.

### Imaging tracers and chemical inhibitors treatment

Cells were kept in culture until reaching 50-80% confluency. The following tracers were then added to pre-warmed cultured medium and incubated on the cells at 37℃ at following time points: DQ-red BSA (5 μg/ml, Invitrogen, Catalog No. D12051) for 1 h 15 min to 6 h 15 min, BSA-Alexa Fluor 647 (10 μg/ml, Invitrogen, Catalog No. A34785) for 1 h 15 min to 6 h 15 min, 40 KDa Dextran-fluorescein (10 μg/ml, Invitrogen, Catalog No. 2306344) for 1 h, wortmannin (0.1 ng/ml, Calbiochem, Catalog No.19545-26-7) for 3 h, L-leucyl-L-leucine methyl ester (LLOMe, 2 mM, Bachem, Catalog No. 400725.0005) for 4 h, CytoFix Red Lysosomal Stain (Cytofix Lyso, 1X, AAT Bioquest, Catalog No. 23210) for 45 min, Magic Red (1x, Abcam, Catalog No. AB270774) for 45 min, SiR-lysosome (SiR Lyso,1 µM, Spirochrome. Catalog No. SC012) for 45 min. Following the incubation, cells were washed with pre-warmed medium. For BSA and dextran tracers, cells were fixed in antigen fix. For lysosomal markers, cells were incubated in PBS-/- and imaging acquisition was performed on Operetta CLS (Perkin Elmer).

### ASOs sequences and synthesis

Single-stranded LNA oligonucleotides were synthesized as described^11^ using standard phosphoramidite chemistry. All sequences have phosphorothioate internucleotide linkages throughout. Production has been done at F.Hoffmann-La Roche, Qiagen or IDT. ASO sequences are annotated following the HELM nomenclature^73^:

HBB splice-switch (SS-ASO) used *in vitro* and *in vivo* : {[LR](A)[sP].[LR]([5meC])[sP].[dR](T)[sP].[dR](T)[sP].[LR](A)[sP].[dR](C)[sP].[dR](C)[sP].[LR](T)[sP].[LR](T

)[sP].[dR](A)[sP].[dR](A)[sP].[LR]([5meC])[sP].[LR]([5meC])[sP].[dR](C)[sP].[LR](A)[sP].[LR](G)}.

MALAT 1 ASO used *in vitro* : {[LR](G)[sP].[LR](A)[sP].[LR](G)[sP].[dR](T)[sP].[dR](T)[sP].[dR](A)[sP].[dR](C)[sP].[dR](T)[sP].[dR](T)[sP].

[dR](G)[sP].[dR](C)[sP].[dR](C)[sP].[dR](A)[sP].[LR](A)[sP].[LR]([5meC])[sP].[LR](T)}

CERS2 ASO used *in vitro* :

{[LR](T)[sP].[LR](T)[sP].[LR](G)[sP].[dR](T)[sP].[dR](T)[sP].[dR](A)[sP].[dR](T)[sP].[dR](T)[sP].[dR](G)[sP].[

dR](A)[sP].[dR](G)[sP].[dR](G)[sP].[dR](A)[sP].[LR](T)[sP].[LR](G)[sP].[LR](G)}

CD81 ASO used *in vitro*:

{[LR](A)[sP].[LR](G)[sP].[dR](T)[sP].[dR](T)[sP].[dR](G)[sP].[dR](A)[sP].[dR](A)[sP].[dR](G)[sP].[dR](G)[sP]

.[dR](C)[sP].[dR](G)[sP].[dR](A)[sP].[dR](C)[sP].[LR](G)[sP].[LR](T)[sP].[LR](G)}

Non-targeting ASO used *in vitro* (CTRL 1 in Supplementary Table 11) :

{[LR]([5meC])[sP].[dR](C)[sP].[LR](A)[sP].[LR](A)[sP].[LR](A)[sP].[dR](T)[sP].[dR](C)[sP].[dR](T)[sP].[dR](T

)[sP].[dR](A)[sP].[dR](T)[sP].[dR](A)[sP].[dR](A)[sP].[dR](T)[sP].[dR](A)[sP].[LR](A)[sP].[LR]([5meC])[sP].[d R](T)[sP].[LR](A)[sP].[LR]([5meC])}

Non-targeting ASO used in Extended data 1a, Supplementary Table 11 (CTRL2) and *in vivo* :

{[LR](G)[sP].[dR](C)[sP].[LR](A)[sP].[dR](A)[sP].[LR](A)[sP].[dR](T)[sP].[LR](T)[sP].[dR](C)[sP].[LR]([5meC])[sP].[dR](T)[sP].[LR](A)[sP].[dR](T)[sP].[LR](T)[sP].[dR](C)[sP].[LR]([5meC])[sP].[dR](C)} .

Fluorescently labeled ASOs of the same sequences ( MALAT1, CERS2 and CTRL1) were synthesized by IDT. Alexa Fluor 647 was attached to the oligo via an NHS esther. The linker used was a C6 Amino modifier, attached to the oligo via a phosphate group.

### siRNA sequences and synthesis

Design, screening and synthesis of ap1m1-GalNAc were performed at Axolabs. The ap1m1-siRNA was chosen by screening dozens of siRNAs bioinformatically with regards to activity and specificity followed by *in vitro* testing in mouse Hepa1-6 cells to find the most potent one. The chosen siRNA was synthesized according to the phosphoramidite technology on a solid support made of controlled pore glass (CPG), employing a Mermade 12 synthesizer (LGC Bioautomation).

Sense strand sequence (5’ to 3’): RNA1{m(G)[sp].m(U)[sp].[fl2r](U)p.m(C)p.[fl2r](G)p.m(U)p.[fl2r](U)p.[fl2r](U)p.[fl2r](C)p.m(A)p.[fl2r](U)p.m(

G)p.[fl2r](U)p.m(G)p.[fl2r](G)p.m(A)p.[fl2r](U)p.m(U)p.[fl2r](A)p.[NHC6]p.[GalNAc3]}$$$$V2.0

Antisense strand sequence (5’ to 3’): RNA1{[vinyl5m](U)[sp].[fl2r](A)[sp].m(A)p.[fl2r](U)p.m(C)p.[fl2r](C)p.m(A)p.[fl2r](C)p.m(A)p.[fl2r](U)p.m(G)p

.m(A)p.m(A)p.[fl2r](A)p.m(C)p.[fl2r](G)p.m(A)p.[fl2r](A)p.m(C)[sp].m(G)[sp].m(C)}$$$$V2.0

Duplex: RNA1{m(G)[sp].m(U)[sp].[fl2r](U)p.m(C)p.[fl2r](G)p.m(U)p.[fl2r](U)p.[fl2r](U)p.[fl2r](C)p.m(A)p.[fl2r](U)p.m(G)p.[fl2r](U)p.m(G)p.[fl2r](G)p.m(A)p.[fl2r](U)p.m(U)p.[fl2r](A)p.[NHC6]p.[GalNAc3]}|RNA2{[vinyl5m](U)[sp]

.[fl2r](A)[sp].m(A)p.[fl2r](U)p.m(C)p.[fl2r](C)p.m(A)p.[fl2r](C)p.m(A)p.[fl2r](U)p.m(G)p.m(A)p.m(A)p.[fl2r](A) p.m(C)p.[fl2r](G)p.m(A)p.[fl2r](A)p.m(C)[sp].m(G)[sp].m(C)}$RNA1,RNA2,2:pair-

56:pair|RNA1,RNA2,5:pair-53:pair|RNA1,RNA2,8:pair-50:pair|RNA1,RNA2,11:pair-

47:pair|RNA1,RNA2,14:pair-44:pair|RNA1,RNA2,17:pair-41:pair|RNA1,RNA2,20:pair-

38:pair|RNA1,RNA2,23:pair-35:pair|RNA1,RNA2,26:pair-32:pair|RNA1,RNA2,29:pair-

29:pair|RNA1,RNA2,32:pair-26:pair|RNA1,RNA2,35:pair-23:pair|RNA1,RNA2,38:pair-

20:pair|RNA1,RNA2,41:pair-17:pair|RNA1,RNA2,44:pair-14:pair|RNA1,RNA2,47:pair-

11:pair|RNA1,RNA2,50:pair-8:pair|RNA1,RNA2,53:pair-5:pair|RNA1,RNA2,56:pair-

2:pair|RNA1,RNA2,59:pair--1:pair|RNA1,RNA2,62:pair—

4:pair$${”RNA1”:{”strandtype”:”ss”},”RNA2”:{”strandtype”:”as”}}$V2.0

The sense strand was assembled on 3’-phthalimidyl-amino-C6-modifier loaded CPG solid support, available from LGC Biosearch Technologies (Petaluma, Catalog No. BG7-1048-B,), with a porosity of 500 Å (86 µmol/g loading). On the other hand, the antisense strand was assembled on 2’-OMe-C loaded CPG solid support, available from LGC Biosearch Technologies (Petaluma, Catalog No. BG7-1030-B), with a porosity of 498 Å (84 µmol/g loading). All 2’-modified RNA phosphoramidites as well as the majority of ancillary reagents were purchased from Merck. Specifically, the following 2’-*O*-methyl phosphoramidites were used: (5’-*O*-dimethoxytrityl-*N6*-(benzoyl)-2’-*O*-methyl-adenosine-3’-*O*-(2-cyanoethyl-*N,N*-diisopropylamino) phosphoramidite, 5’-*O*-dimethoxytrityl-*N4*-(acetyl)-2’-*O*-methyl-cytidine-3’-*O*-(2-cyanoethyl-*N,N*-diisopropylamino) phosphoramidite, (5’-*O*-dimethoxytrityl-*N2*-(isobutyryl)-2’-*O*-methyl-guanosine-3’-*O*-(2-cyanoethyl-*N,N*-diisopropylamino) phosphoramidite and 5’-*O*-dimethoxytrityl-2’-*O*-methyl-uridine-3’-*O*-(2-cyanoethyl-*N,N*-diisopropylamino) phosphoramidite. The 2’-deoxy-2’-fluoro phosphoramidites carried the same protecting groups as the 2’-*O*-methyl RNA amidites. The (vinu)-nucleotide unit incorporated at the 5’-end of the antisense strand was derived from 5’-*bis*POM-(*E*)-vinyl phosphonate-2’-*O*-methyl-uridine phosphoramidite procured from LGC Biosearch Technologies (Petaluma, Catalog no. LK2562). All amidites were dissolved in anhydrous acetonitrile (100 mM) and molecular sieves (3 Å) were added. 5-Ethylthiotetrazole (ETT, 500 mM in acetonitrile) was used as an activator solution. Coupling times were 6 min. In order to introduce phosphorothioate linkages, a 100 mM solution of 3-amino-1,2,4-dithiazole-5-thione (obtained from TCI Chemicals) dissolved in ACN-pyridine (2:3 *v*/*v*) was employed as sulfurizing agent. TCA (3% in dichloromethane) was used as a deblocking agent. The other ancillary reagents used were as follows: Iodine-oxidizer (50 mM I2 in pyridine-H2O (9:1)), Cap A (acetic anhydride in THF (9.1:90.9 *v/v*)) and Cap B (THF, *N*-methylimidazole and pyridine (8:1:1 *v/v/v*)) as capping agents. The sense strand was synthesized with removal of the final DMT protecting group (“DMT-Off”).

After finalization of the solid phase synthesis, the sense strand immobilized CPG solid support was treated with AMA (1:1 *(v/v)* mixture of concentrated aqueous ammonia and 40% aqueous methylamine). The antisense strand immobilized CPG solid support, on the other hand, was treated with 3% DEA/NH3 (3:97 *(v/v)* mixture of 99.5% diethylamine and concentrated aqueous ammonia (solution (30%, puriss) in water), to ensure cleavage from the solid support and quantitative removal of all the protecting groups. The oligonucleotides were finally precipitated overnight in the freezer using 3 M NaOAc (pH 5.2) in ethanol. Pellets were reconstituted in RP-HPLC Buffer A and were purified by preparative RP-HPLC using an XBridge BEH C18 (19×50 mm, OBD, 5 µm particle) prep-column (WATERS GmbH) on an ÄKTA Pure 150 system (Cytiva). Buffer A was 100 mM triethylammonium acetate (TEAA, pH 7) in water and buffer B was a formulation of buffer A in 95% acetonitrile-water. UV traces at 260 and 280 nm were recorded to monitor product elution. A gradient from 0% B to 100% B was applied. Appropriate fractions were pooled and precipitated overnight in the freezer using 3 M NaOAc (pH 5.2) in ethanol. Pellets were reconstituted in water and were quantified by measuring the UV absorption at 260 nm. Oligonucleotide concentrations were calculated based on their theoretical extinction coefficients calculated by the ‘nearest neighbor’ method.

The peracetylated free-acid derivative of the triantennary GalNAc cluster (GalNAc3-free acid) was prepared pursuing disclosures from^74^. For GalNAc conjugations to the free amine on the 3’-terminal of the sense strand, the free acid derivative was activated by conversion to the NHS-ester using *N*-hydroxysuccinimide (NHS, Sigma-Aldrich, Catalog No.130672) and *N,N*′-diisopropylcarbodiimide (DIC, Sigma-Aldrich, catalog No. D125407) and finally was conjugated to the hexylamino-linker at the 3’-end of the sense strand. After optimal conversion/conjugation, the reaction mixture was diluted with water, filtered through a syringe filter and then purified by preparative RP-HPLC using an XBridge BEH C18 column using the same protocol as before. Aqueous solutions of the GalNAc3 (acetyl-protected)-conjugated sense strand were treated with aqueous ammonia solution (30%, puriss) and incubated with shaking to ensure deacetylation. Thereafter, ammonia was removed by evaporation in a vacuum concentrator, followed by precipitation overnight in the freezer using 3 M NaOAc (pH 5.2) in ethanol.

The purified GalNAc-conjugated sense strand and the antisense strand were quantified by UV absorption at 260 nm and were assessed for identity (±0.05% of calculated molecular weight (by ESI-MS) and purity (single strand purity > 85%, as per integration of the UV signal of the analytical RP-HPLC trace).

The purified complementary strands were then mixed in an equimolar ratio to yield 15.5 mg of the desired ap1m1-siRNA in purified water. The duplex solutions were placed into a water bath at 70 °C followed by subsequent cooling to room temperature. The resultant siRNA-duplex was characterized by size-exclusion chromatography (SEC) and exhibited a duplex purity > 90%, as per integration of the UV signal of the analytical SEC trace.

The GalNAc ahsa1 (aha-1) siRNA was designed and synthesized at F. Hoffmann-La Roche using standard phosphoramidite chemistry followed by a post-conjugation of the tri-GalNAc to the C6 amino linker (Nair 2014). In the below description of the chemical modifications of the siRNA, the phosphorothioate modifications have been indicated with a “PS ’’ otherwise, the internucloside linkage is synthesized as a phosphodiester “PO”. The final duplex was after annealing characterized w. a 93% purity and a calculated MW of 14757.16 g/mol (found MW of 14755.9 g/mol).

Sense strand sequence w. 3’ aminohexyl tri-GalNAc (5’ to 3’): UCUCGUGGCCUUAAUGAAA-C6-amino-(GalNAc)3. Modification pattern of sense strand from 5’ to 3’: 5’ - [2’F]-PS-[2’OMe]-PS-[2’F]-[2’OMe]-[2’F]-[2’OMe]-[2’F]-[2’OMe]-[2’F]-[2’OMe]-[2’F]-[2’OMe]-[2’F]-[2’OMe]-[2’F]-[2’OMe]-[2’F]-[2’OMe]-PS-[2’F]-PS-[C6-amino-linker]-[Gal-NAc]3 3’

Antisense strand sequence (5’ to 3’): UUUCAUUAAGGCCACGAGAUU

Modification pattern of antisense strand from 5’ to 3’: 5’ - [2’OMe]-PS-[2’F]-PS-[2’OMe]-[2’F]-[2’OMe]-[2’F]-[2’OMe]-[2’F]-[2’OMe]-[2’F]-[2’OMe]-[2’F]-[2’OMe]-[2’F]-[2’OMe]-[2’F]-[2’OMe]-[2’F] - [2’OMe] –PS-[2’OMe] –PS-[2’OMe]-3’

### Rodent husbandry

Mice were group-housed in a 12-h light/dark cycle (light between 07:00 and 19:00) in a temperature-controlled room (21.1 ± 1.1℃) with free access to water and food in the animal facility of the F. Hoffmann-La Roche site in Basel. All animals in this study are sourced from Charles River (Germany) and are from the following line: C57BL/6J-Gt(Rosa)26Sor<em1(CAG-mKate2,-EGFP*)Gne, background strain C57BL/6, homozygous knock in. This line has been generated by^17^. Both male and female adult mice were used (12-18 weeks old at sacrifice time). All experiments were done under the swiss license # 35726 (National No.) and #3196 (Basel Cantonal No.). The group allocation was performed such as to minimize the effect of age, weight, sex, and housing arrangements.

### Administration of substances and sacrifice for *in vivo* experiments

Both GalNAc-siRNA (15 mg/kg) and ASOs (10 mg/kg) were administered sub-cutaneously (see sequences above). SiRNA were administered at day 0, ASOs at day 15, and mice were sacrificed at day 18. On the day of sacrifice, mice were injected intraperitoneally with pentobarbital (150 mg/kg). Upon reaching deep anesthesia, animals underwent transcardiac perfusion with PBS, followed by sacrifice of the liver and kidney samples. Tissues were stored in Precellys tubes (Bertin Technologies, Catalog No. P000911-LYSK1-A) in dry ice, and then at -80 ℃.

### Extraction of RNA and qRT-PCR for *in vivo* samples

RNA extraction was performed with the Precellys Tissue total RNA extraction kit (Bertin Technologies, Catalog No. D05711) according to the manufacturer protocol. Briefly, snap frozen tissue were lysed in 700 µl of RNA lysis buffer provided in the PrecellysTissue total RNA extraction kit and mechanically dissociated with the Precellys Evolution Tissue homogeniser (Bertin Technologies, 2 cycles of 20 seconds 6000x RPM with a 10 seconds break). Samples were centrifuged at 10000x g for 5 min and the clear supernatant was transferred into a fresh tube containing an equal volume of 70% ethanol. After thorough mixing, samples were transferred, by batches of 700 µl, into RNA mini columns and centrifuged at 10000x g for 1 min. Columns were washed with Wash buffers I and II, with centrifugation steps of 30 sec in between. After drying the membrane with a 2 min centrifugation at 10000x g, RNA was eluted in 100 µl of RNase free water (Thermo Fisher, Catalog No. AM9937, centrifugation at 10000x g, 2 min). RNA yield and quality was assessed using a Nanodrop 8000 (Thermo Fisher).

Reverse transcription was performed using components of the Cells-to-CT Bulk RT Reagents (Thermo Fisher, 4391852C) and following manufacturer’s instructions. Briefly, each reaction contained approx. 10 ng of RNA freshly extracted, RT buffer (1X), and RT enzyme mix (1X), in a final volume of RNase free water (Thermo Fisher, Catalog No. AM9937) of 50 µl. The thermocycler program was the following: 10 minutes at 42℃, 1 hour at 37℃ then hold at 4℃. QRT-PCR was performed using the TaqPath™ 1-Step Multiplex Master Mix (No ROX) (Thermo Fisher, Catalog No. A28523), and the primers targeting ap1m1, ahsa1, and gapdh (all Thermo Fisher with corresponding Catalog Nos.; ap1m1 ; Mm00475912_m1, ahsa1; Mm01296842_m1, gapdh; Mm9999915_g1). Each reaction contains 1µl of cDNA, 2.5 µl of Taqpath master mix (Thermo Fisher, Catalog No. A15297), and 0.16 µl of each primer (stock 60x) in a final volume of 10 µL in RNase free water (Thermo Fisher, Catalog No. AM9937). The thermocycler Quantstudio Flex 12k Flex (Thermo Fisher) was used using a 40 cycles program.

### Protein extraction and Western blot from *in vitro* and *in vivo* samples

For *in vitro* samples, media was removed and cells washed twice with PBS before being lysed on plate in Pierce/Thermo RIPA buffer (Thermo Fisher, Catalog No. 89900) complemented with protease inhibitor cocktail (Thermo Fisher, Catalog No. 1861281). Samples were sonicated for 20 seconds, and kept on ice for 20 min. Samples were then centrifuged at high speed and only the supernatant was kept and stored at -20℃ for long term storage. Proteins were quantified using the MicroBCA kit (Thermo Fisher, Catalog No. 23235) following the manufacturer’s instructions. Briefly, samples (1:200 dilution) were incubated at 37℃ for 1 h with a solution containing 50% Reagent A, 48% Reagent B and 2% reagent C. Absorbance at 562 nm was measured with an Enspire 2300 reader (PerkinElmer), and the concentration was determined according to the measured albumin standards. Ten μg of protein were separated in 4–12% BIS-TRIS gels and transferred to nitrocellulose or PVDF membranes via iBlot (Thermo Fisher). Membranes were blocked in the Intercept blocking buffer (Licor, Catalog No. 927-70001). The following antibodies were used at 1:1000 with a Intercept blocking buffer (PBS, Licor, Catalog No. 927-70001) containing 0.2% Tween 20 overnight at 4℃: AP1M1 (Thermo Fisher, Catalog No. PA5-98842) and TBC1D23 (Proteintech, Catalog No. 17002-1-AP). The next day, membranes were washed and incubated in the Intercept blocking buffer containing 1:10000 of antibody against actin (Abcam, Catalog No. ABCAAB6276-50) for 1 h at room temperature. Following washes in PBS with 0.1% Tween, the species-relevant secondary antibodies (LI-COR, Catalog Nos. 926-32213, 926-68072) were used at 1:20000 in blocking solution containing 0.2% Tween. Protein was visualized using an Odyssey CLx imager (Licor).

For *in vivo* samples, protein extraction was performed with the reagents from the GFP Elisa kit (Abcam, Catalog No. ab171581), following the manufacturer’s protocol. Briefly, snap frozen tissues were lysed in 300 µl of a solution containing Cell Extraction Buffer PTR (Abcam, Catalog No. ab193970, 1x), enhancer solution (provided in the kit, 1x), protease inhibitor cocktail (Thermo Fisher, Catalog No. 1861281, 1x), and mechanically dissociated with the Precellys Evolution Tissue homogenizer (see above). Samples were incubated on ice for 20 min and then centrifuged at 10000x g at 4℃ for 20 min. The clear supernatant containing the proteins was transferred to a clean tube and kept frozen until the day of the protein quantification. Proteins were quantified using the MicroBCA kit (Thermo Fisher, Catalog No. 23235) as described above. Western blot was performed with 10 µg of protein, according to the protocol described above, with the following antibodies: AP1M1 1:1000 overnight 4℃ (Proteintech, Catalog No. 12112-1-ap) and albumin 1:10000 1 h at room temperature (Bethyl laboratories, Catalog No. a90-134a).

### GFP ELISA

GFP Elisa was performed on *in vivo* samples, following protein extraction as described above, with a GFP Elisa kit according to the manufacturer protocol (Abcam, Catalog No. ab171581). Briefly, protein samples were incubated with a solution containing both the Capture and the Detector antibodies (1X) in antibody diluent solution (1X), in the provided ELISA plate, for 1 h on a shaker (500x RPM). Following the incubation with the antibodies, the plates were washed 3 times with a wash buffer (provided in the kit, 1X) and dried by flipping them on an absorbing towel. Hundred µl of the TMB development solution was added to the plate and the kinetic colorimetric measurement in Enspire 2300 reader at 600 nm (Perkin Elmer) was started. The amount of GFP protein was determined according to the provided GFP standard curve of standards.

## Data and Statistical analysis

For all data and statistical analysis that was not explained in further details above, prism 7 (GraphPad) was used to create charts and perform statistical analyses. Statistical analysis is reported in the figure legends. The tests used included unpaired, two-tailed Student’s t-test, one way ANOVA, and 2-way ANOVA. As post-tests Dunnet, Tukey and false discovery rate (FDR) were used. For all bar graphs, data are represented as mean ± SD, mean ± SEM, or as single points and median or mean. P-values < 0.05 were considered significant, unless stated otherwise.

## Source Data

The normalized read counts, and summarized per gene values for all CRISPR screens are provided as Supplementary Tables 2 and 3. Unprocessed Western Blots are available in source data. RNA-seq expressions are provided as Supplementary Tables 10, 11 and 12. FASTQ files for the RNA-seq experiments are deposited at GEO under accession numbers:GSE280430 and GSE280432

## Supporting information

uncropped blots

## Acknowledgments

We would like to thank Spyros Panfilos, Martine Kapps, Nicole Soder, Silke Zimmermann and Julia Hesselmann for technical assistance. We would like to thank Lars Jønson for the initial discussion on the design of the splice constructs and Helene Gylling, Hendrik Knoetgen, Mark Burcin and Claas Aiko Meyer (F. Hoffman-La Roche) for initial discussions on the project. We would like to thank Ted Lau for the help with DNA isolation, Colin Watanabe for the design of the CRISPR libraries (all Genentech Inc.). We would like to thank the Genentech DNA/RNA sequencing (Jessica Lund), virus (Jessy Sheng, Yuxin Liang and Honglin Chen) and animal (Soren Warming) core facilities for helping with the NGS and viral production. We would like to thank the Genentech FACS core (Ck Poon) and the FACS core of F. Hoffmann-La Roche. We would like to thank Erik Funder for providing the siRNA-GalNAc against ahsa1. Electron

microscopy was performed by the BioEM facility of the Biozentrum (University of Basel) and we would like to thank Cinzia Tiberi Schmidt for her support. Funding for this study was supplied by F. Hoffmann-La Roche and Genentech Inc.

