## Supplementary figures and images for "A CRISPR/Cas9 screen reveals proteins at the endosome-Golgi interface that modulate cellular ASO activity"

### Extended data 1

Extended data 1. Malong et al.

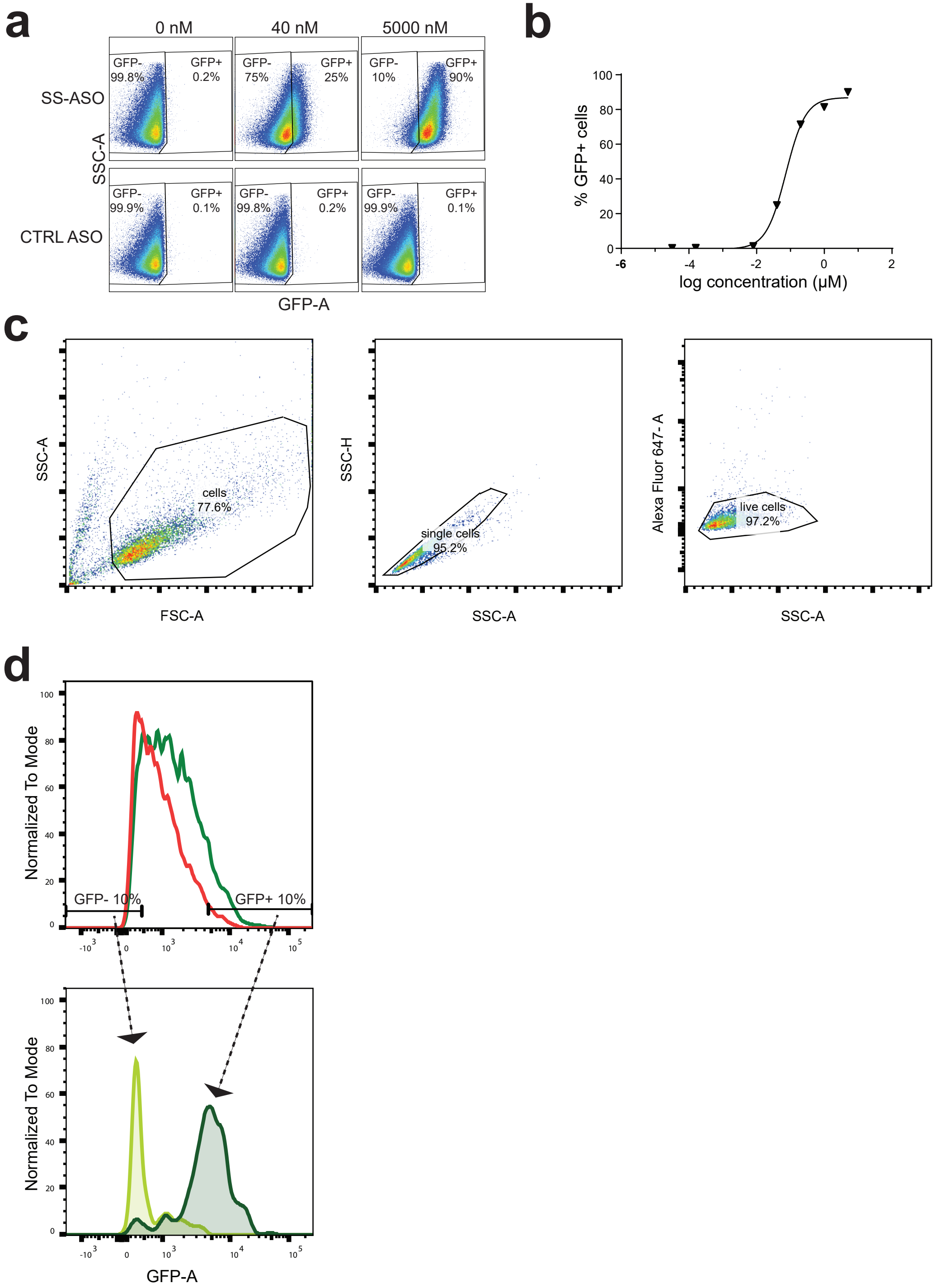

### Extended data 2

Extended data 2. Malong et al.

**a**

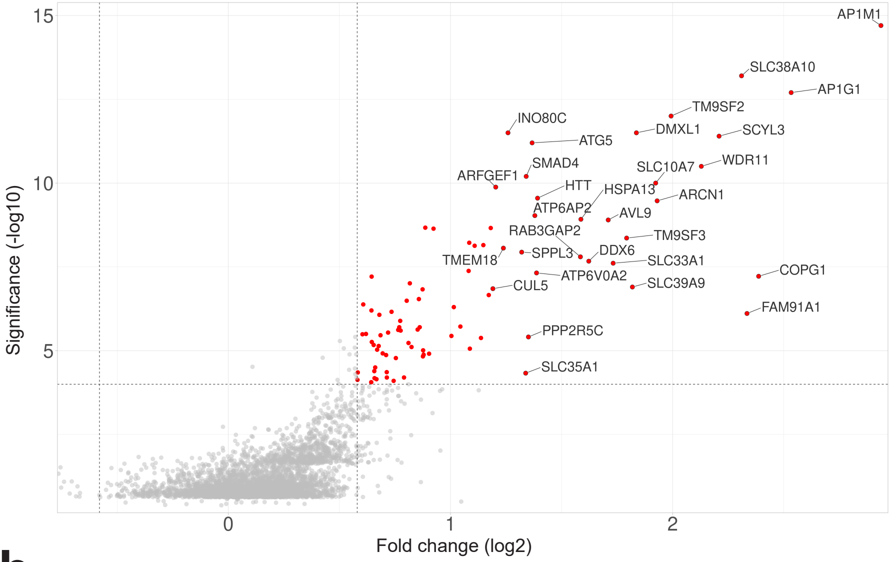


**b**

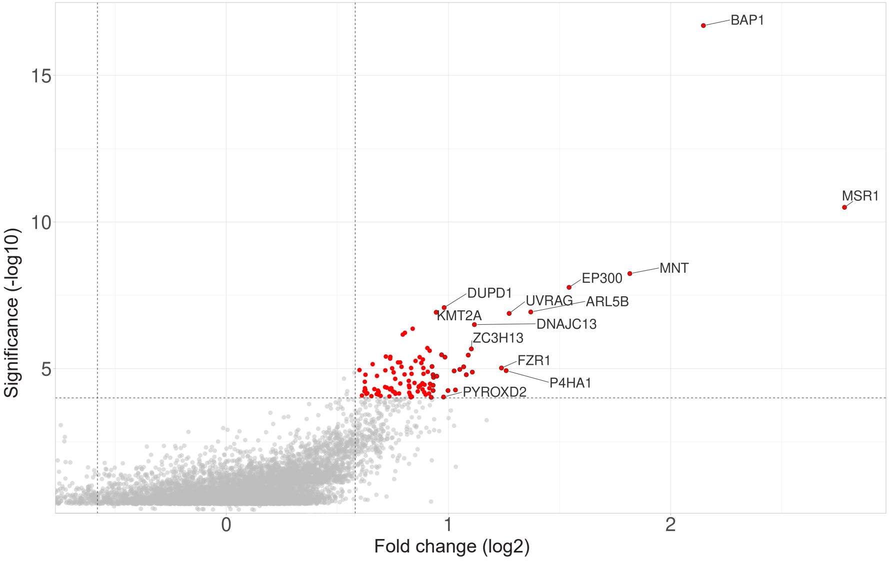

### Extended data 5

Extended data 5. Malong et al.

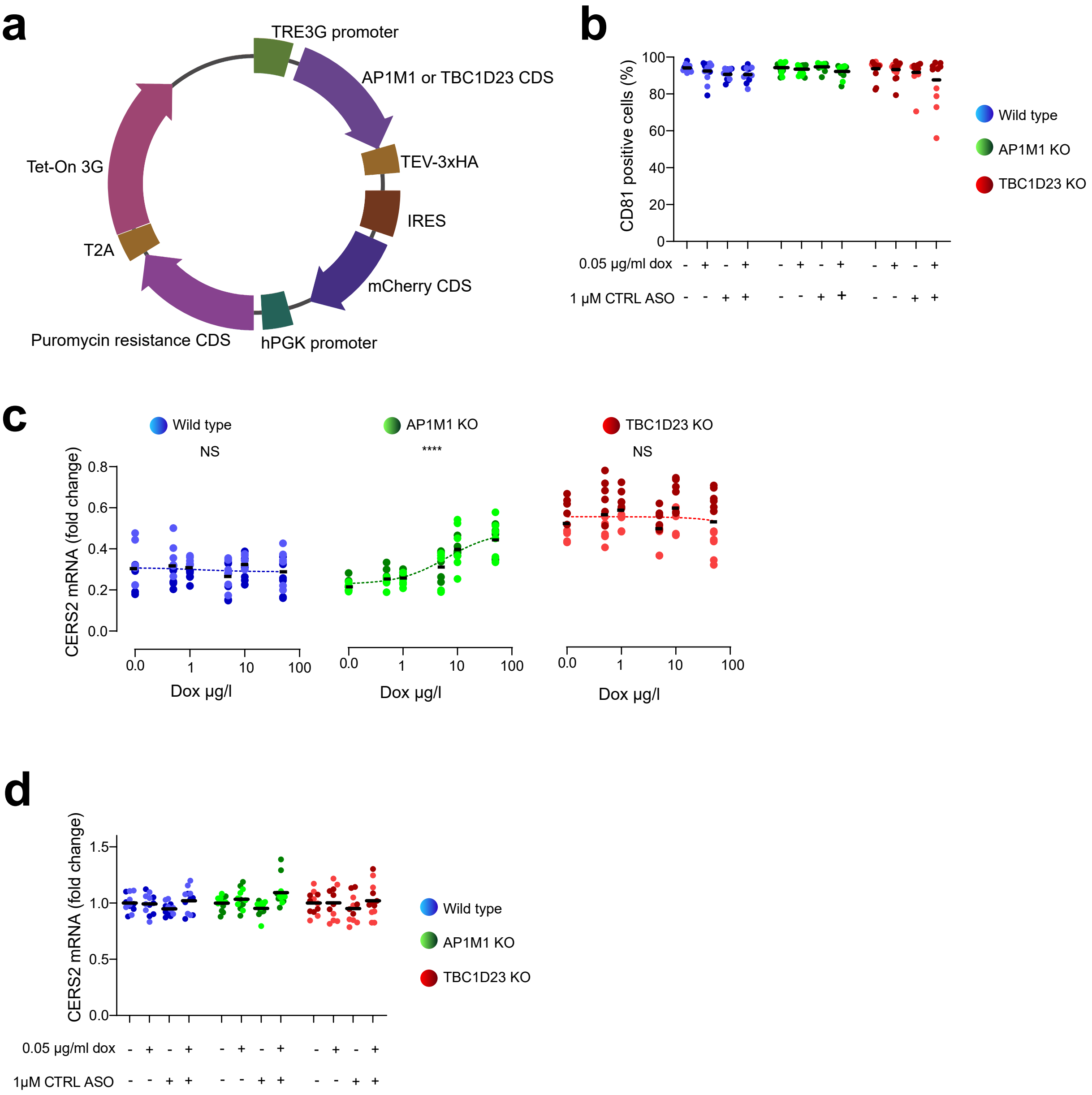

### Extended data 6

Extended data 6. Malong et al.

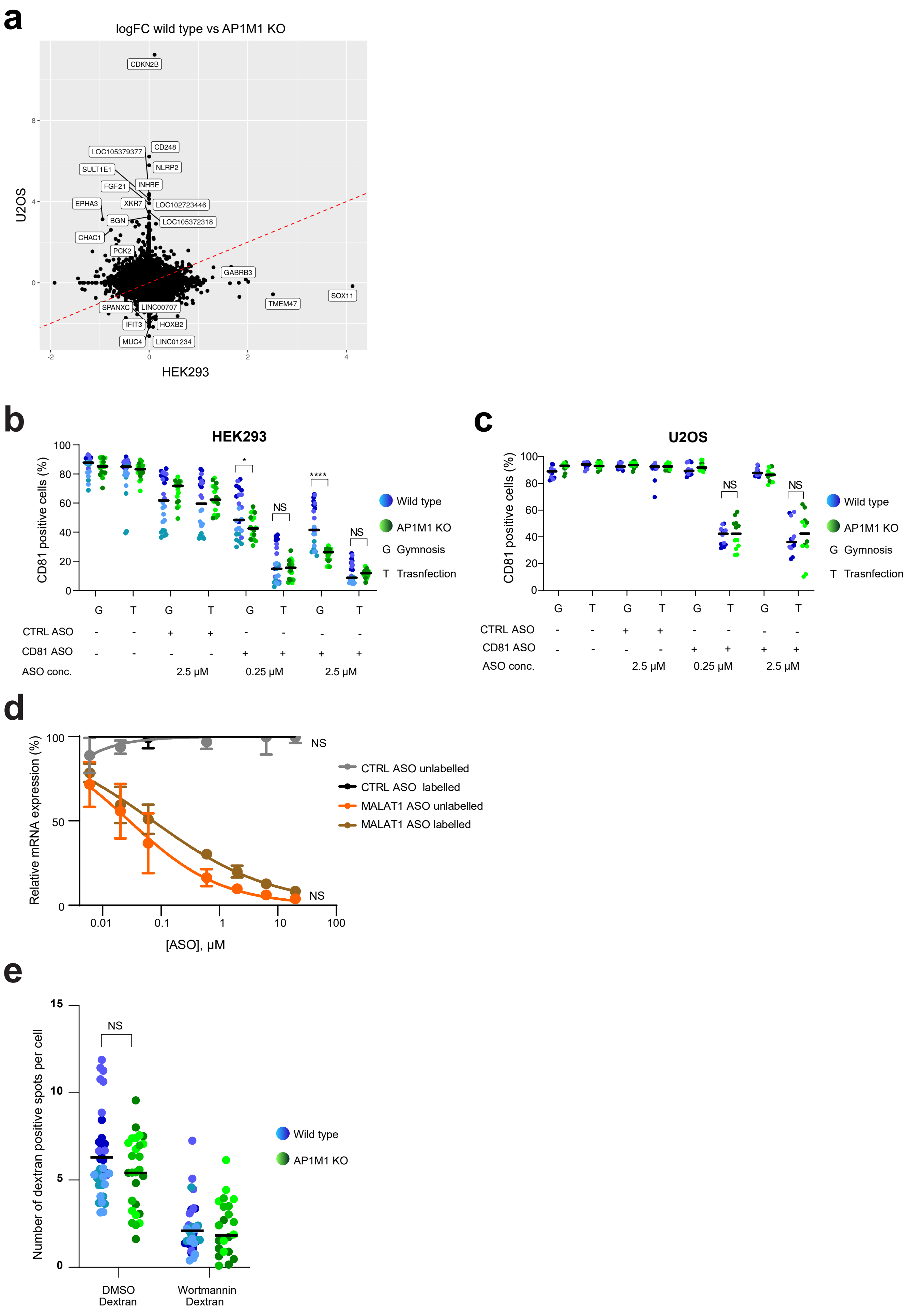

### Extended data 7

Extended data 7. Malong et al.

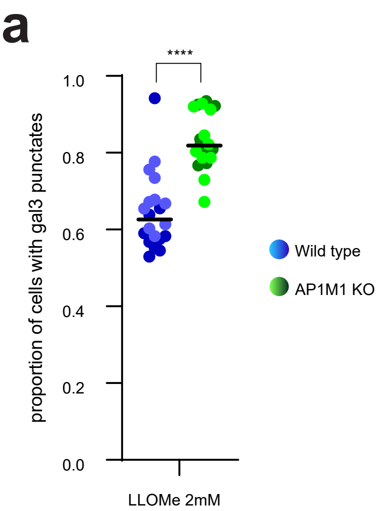


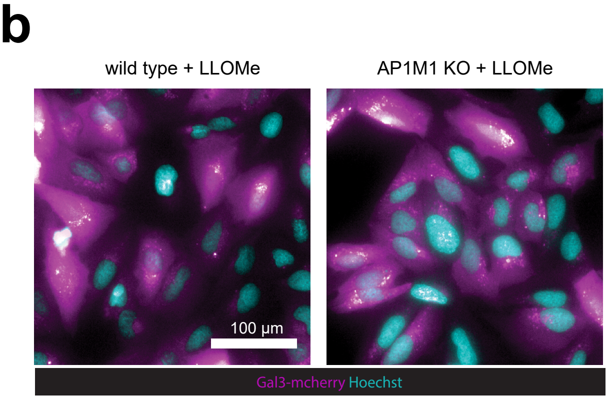

### Extended data 8

Extended data 8. Malong et al

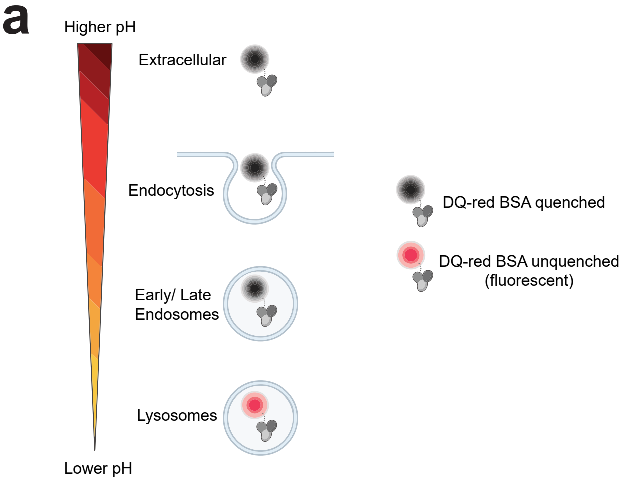


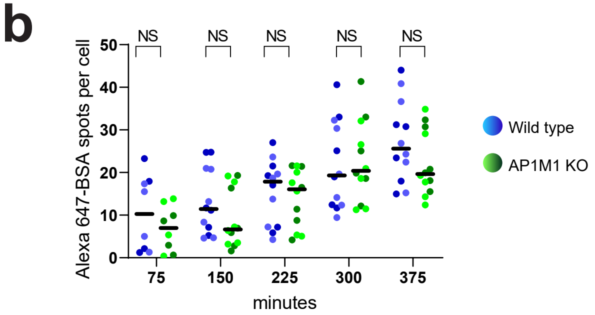

### Extended data 9

Extended data 9. Malong et al.

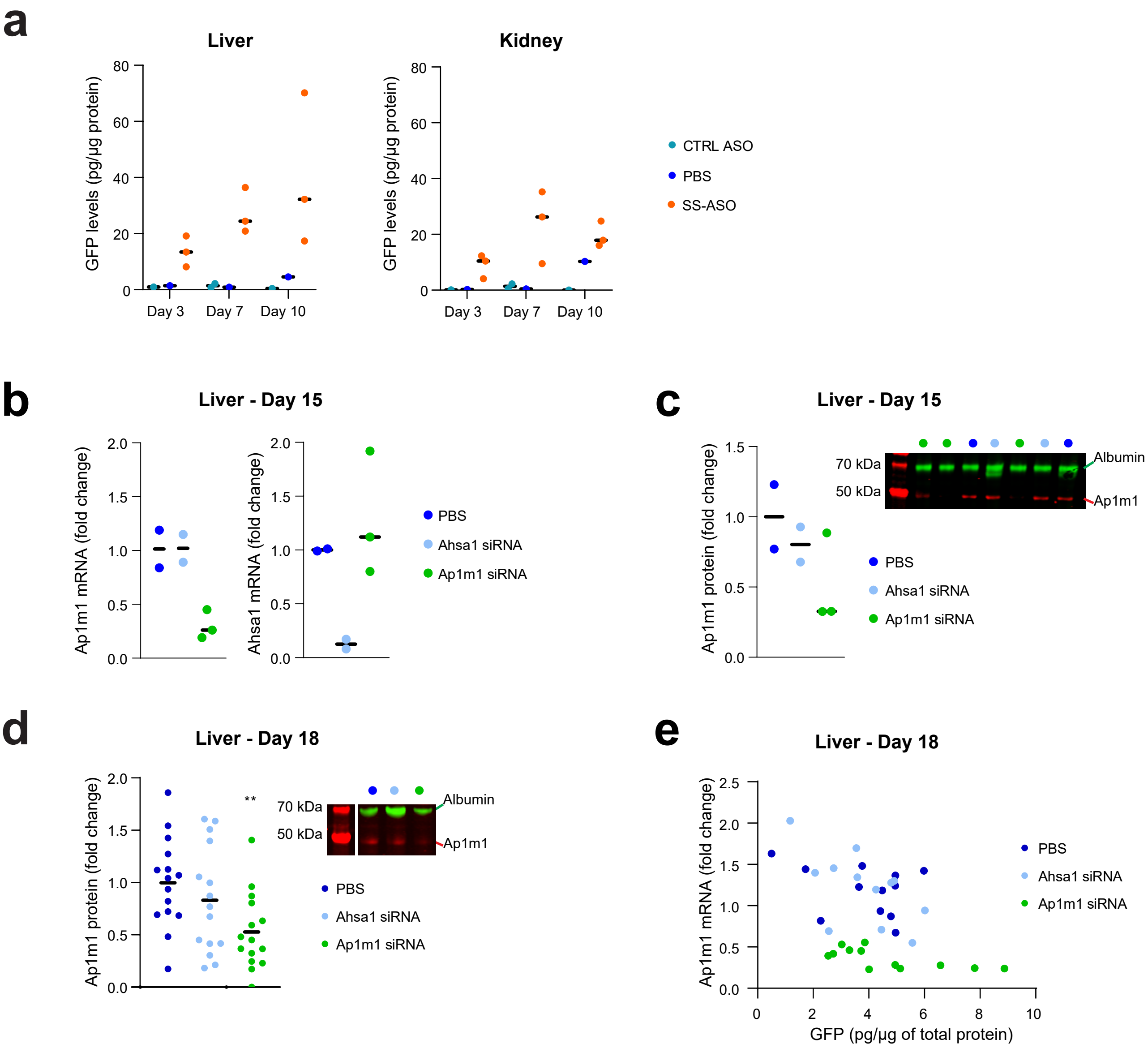

### uncropped blots

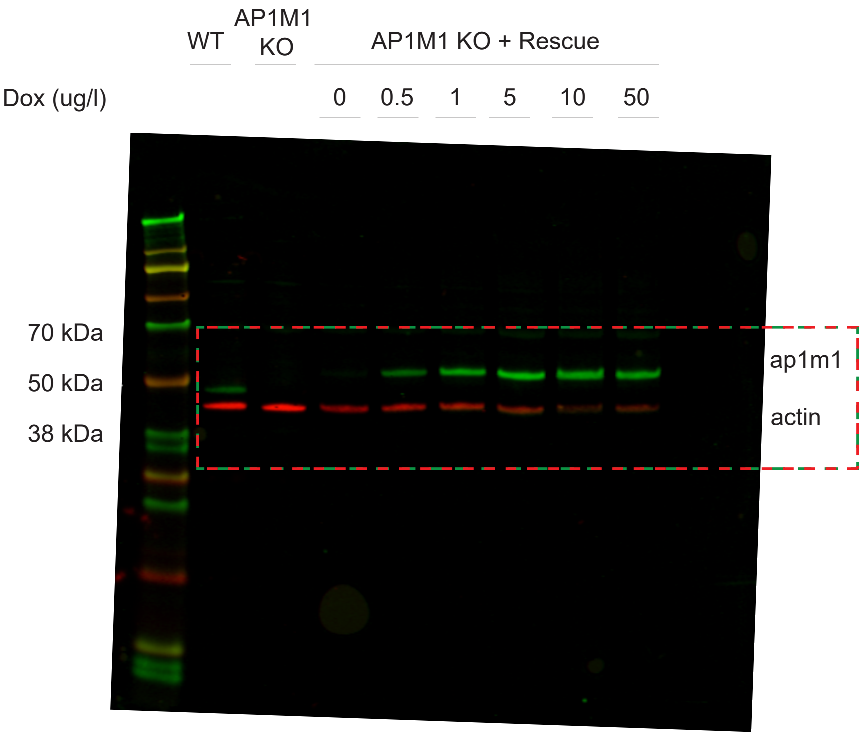

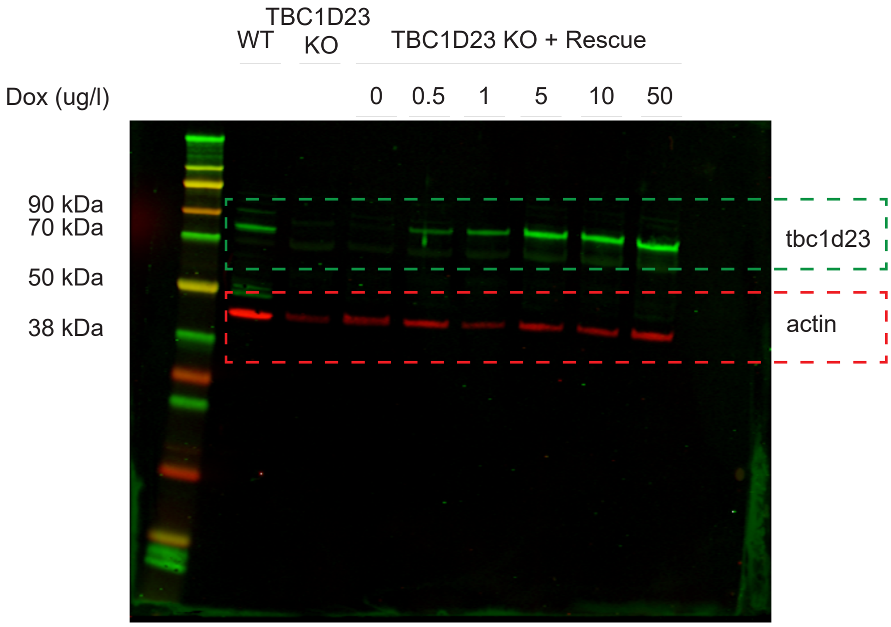

Extended Data 3c

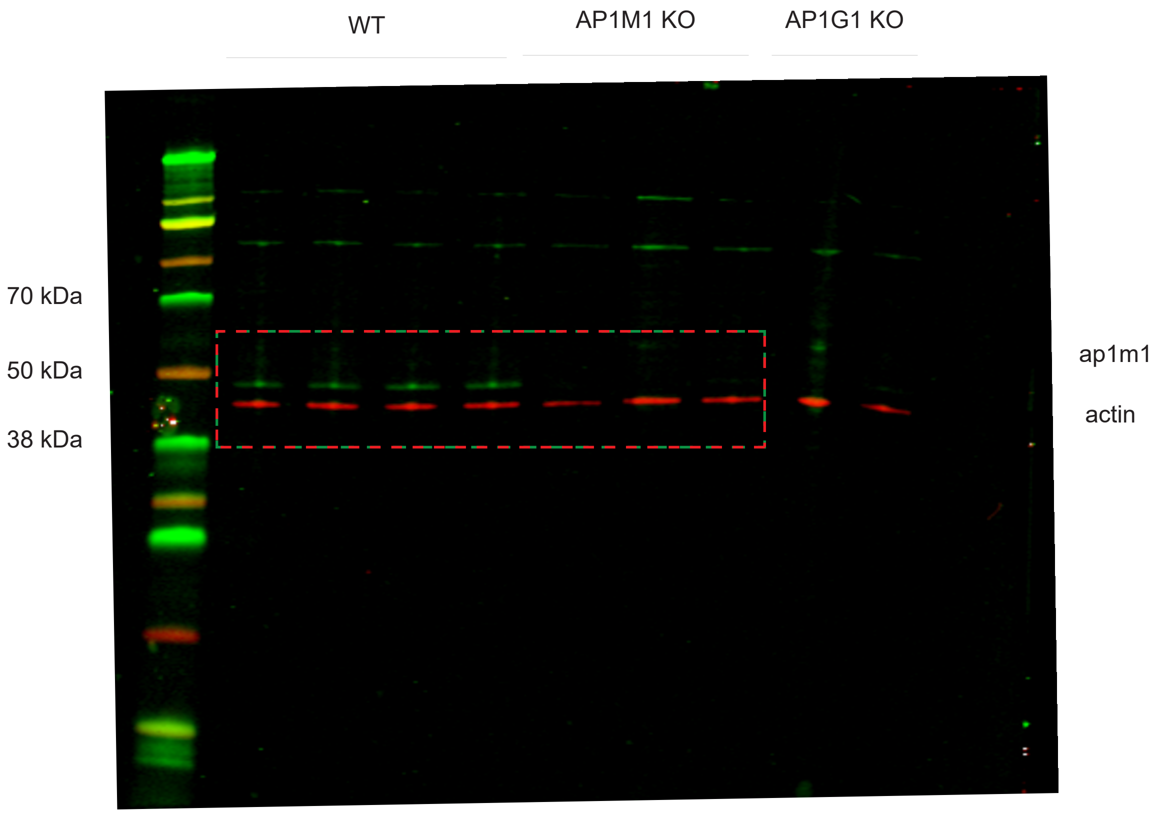


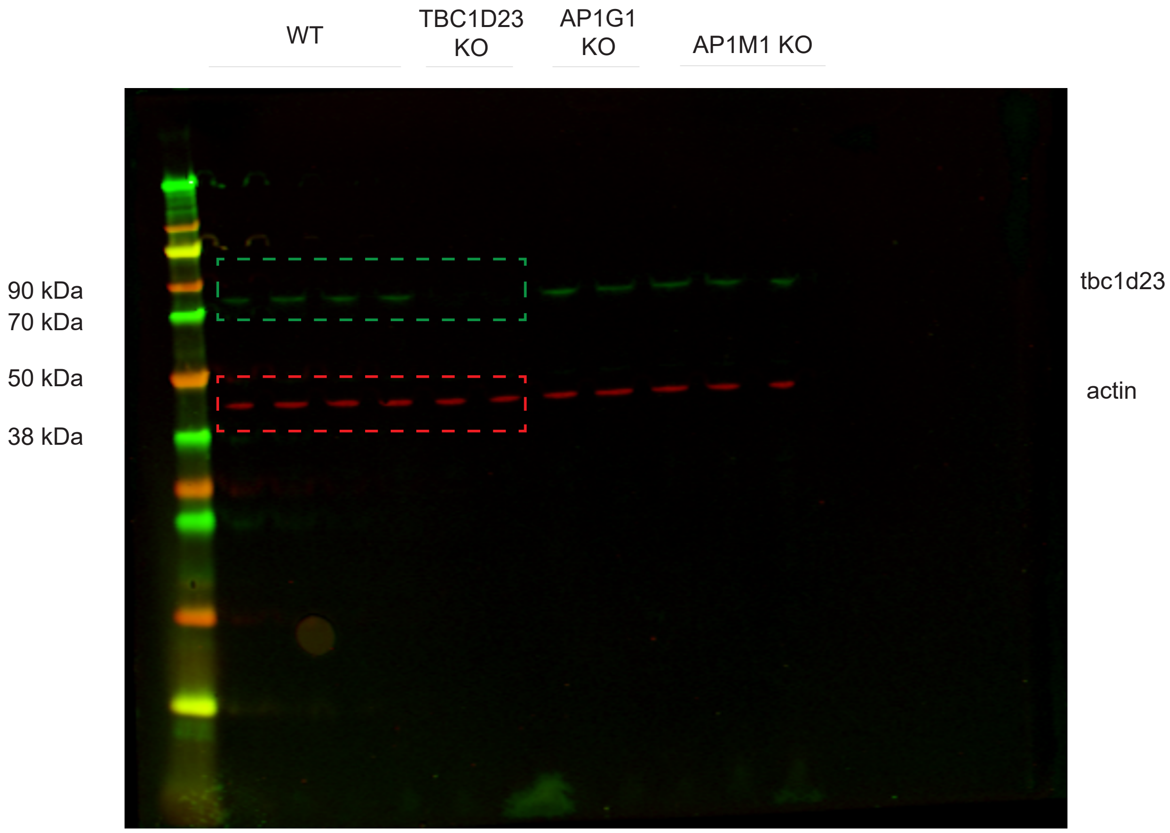

Extended Data 4c

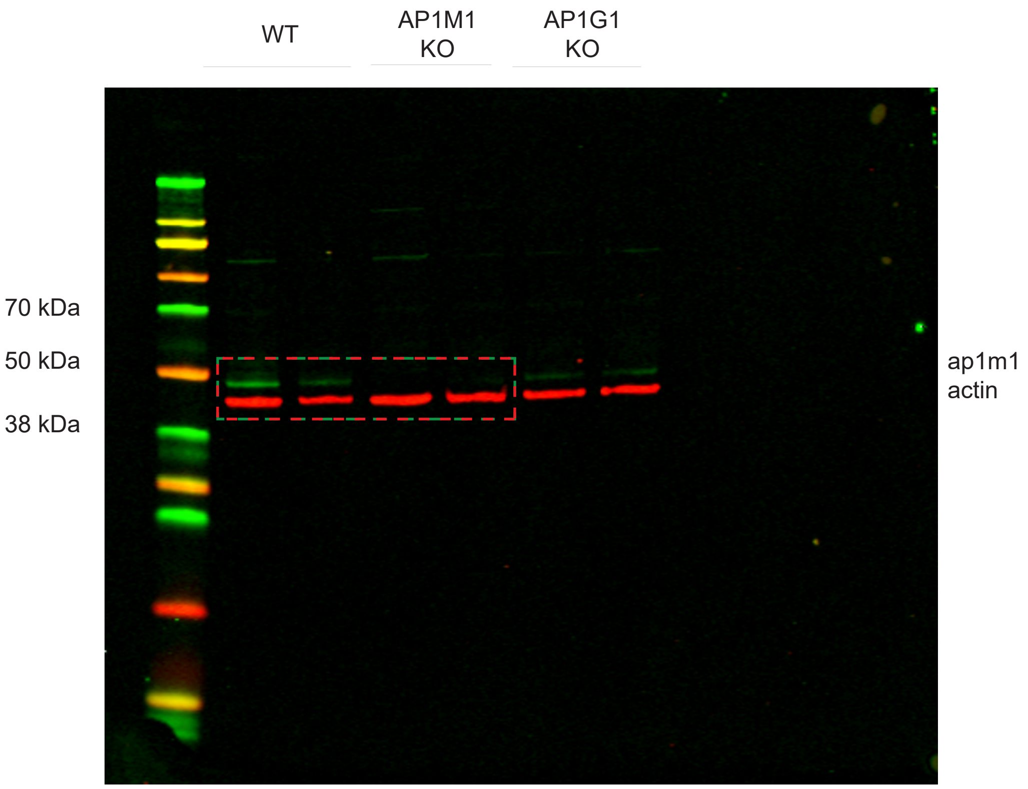


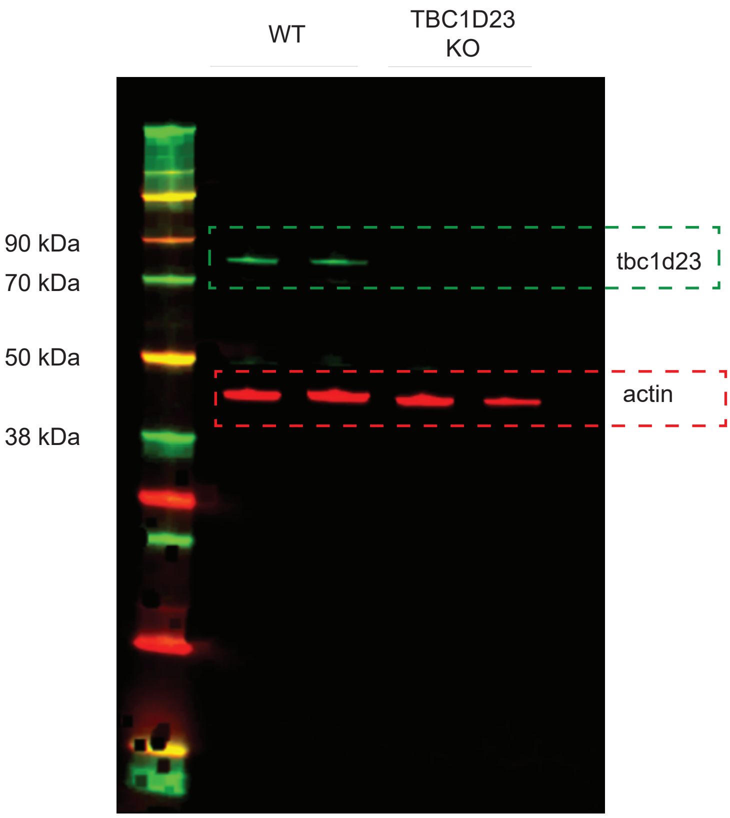

Extended Data 9c

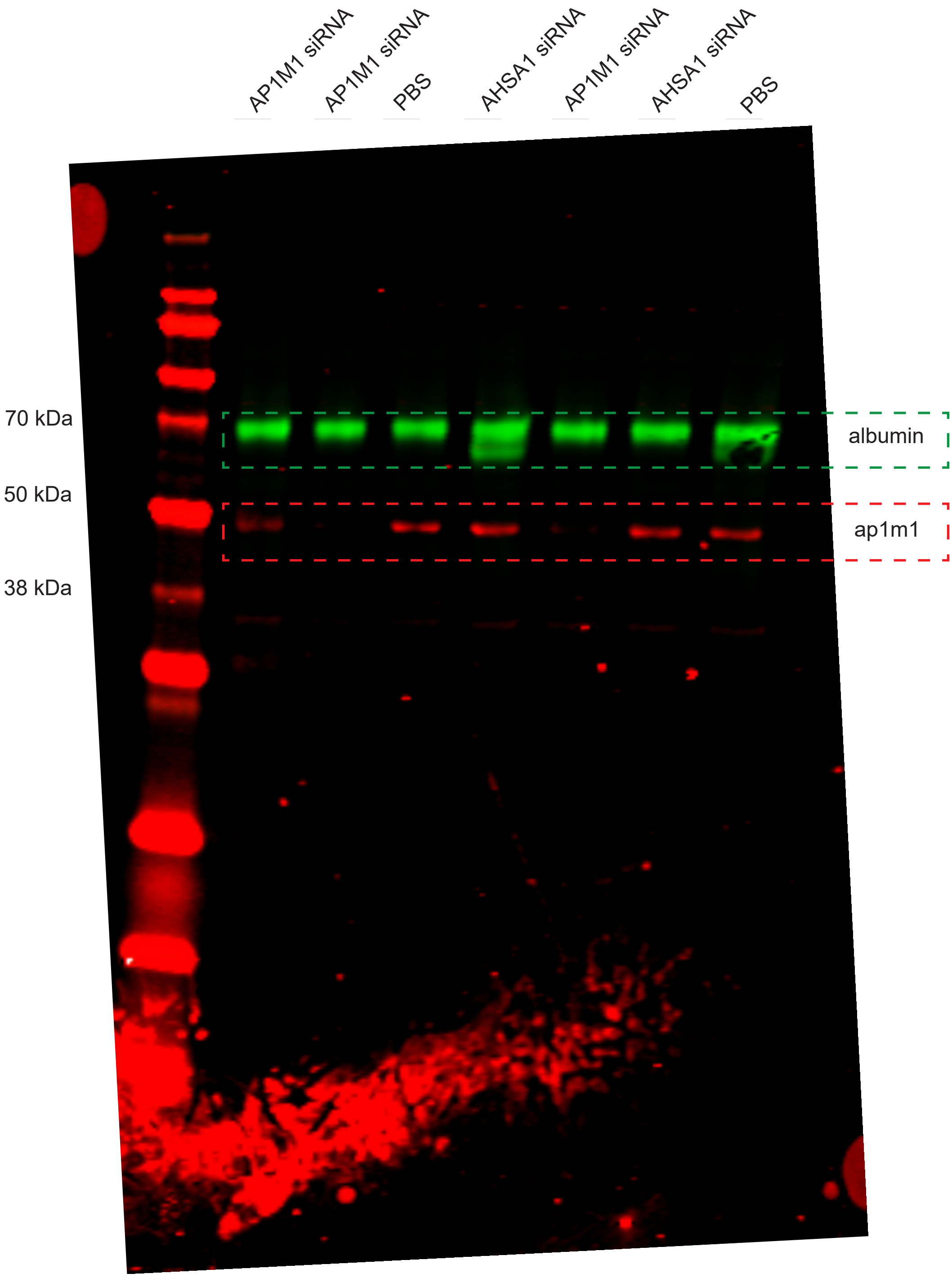

Extended Data 9d

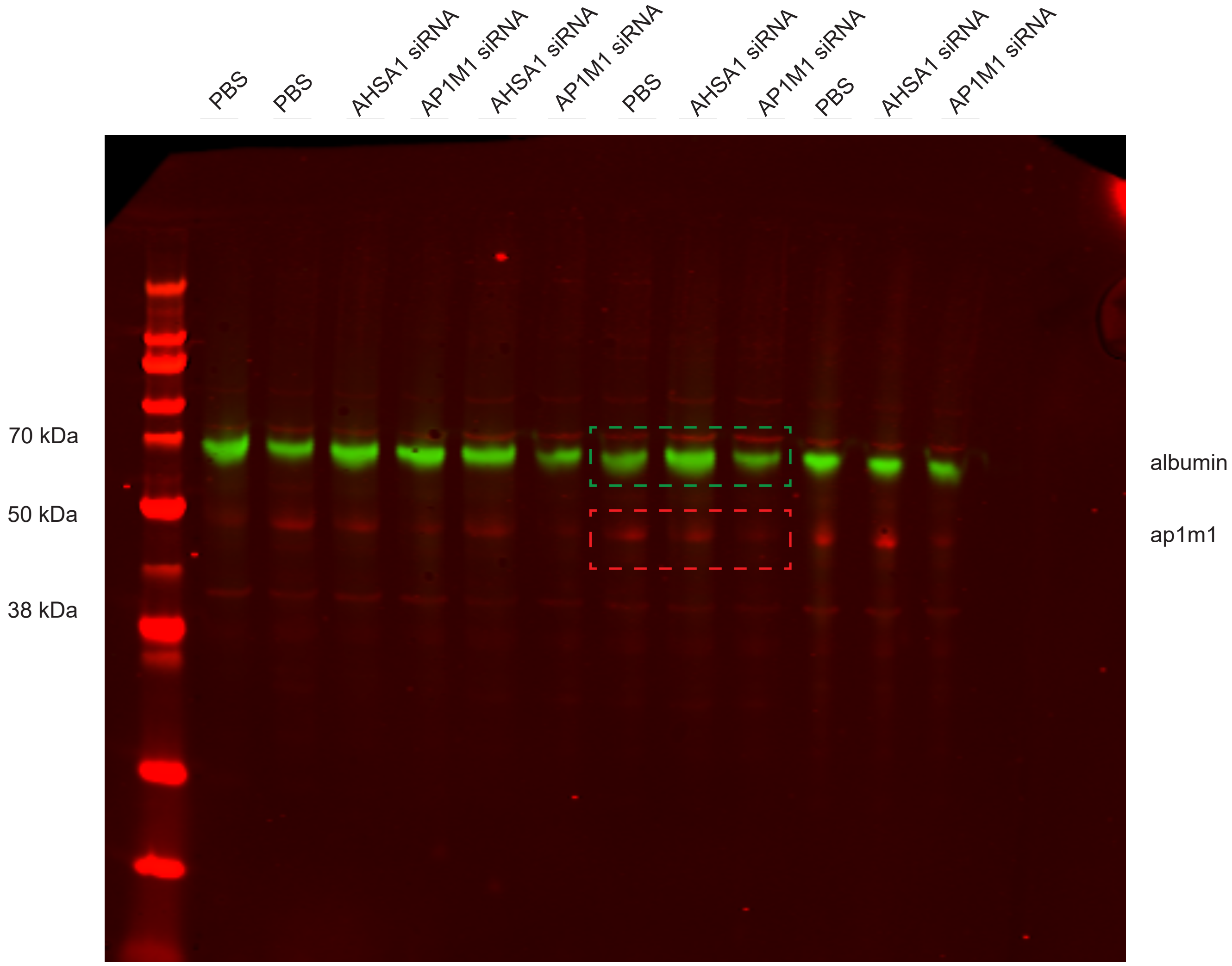
