## Extended data 3 for "A CRISPR/Cas9 screen reveals proteins at the endosome-Golgi interface that modulate cellular ASO activity"

Extended data 3. Malong et al.

**a**

**HEK293**

| Editing frequency of modified CRISPResso2 reads | |
|---|---|
| AP1M1 KO (1) | 98.33% |
| AP1M1 KO (2) | 99.87% |
| AP1M1 KO (3) | 99.96% |
| TBC1D23 KO (1) | 99.51% |
| TBC1D23 KO (2) | 97.94% |

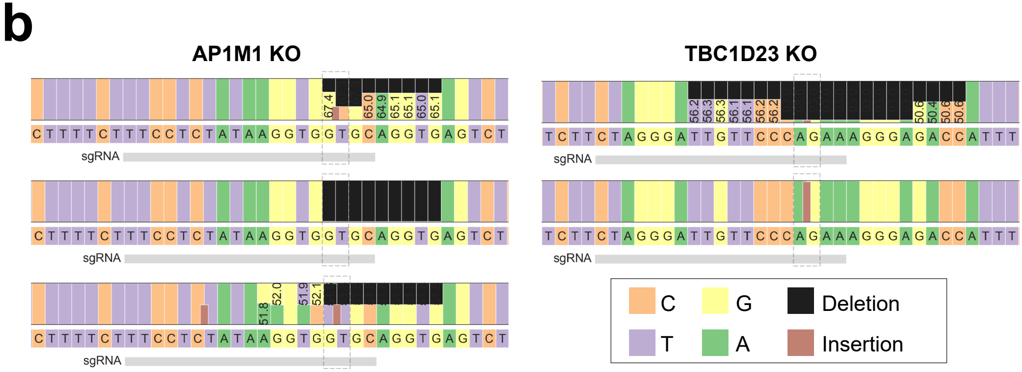


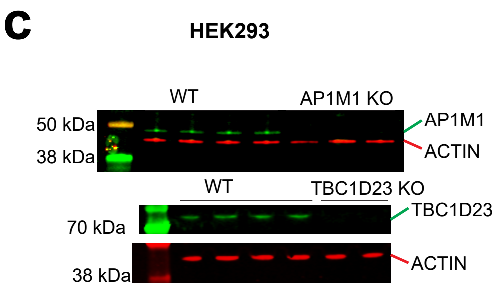


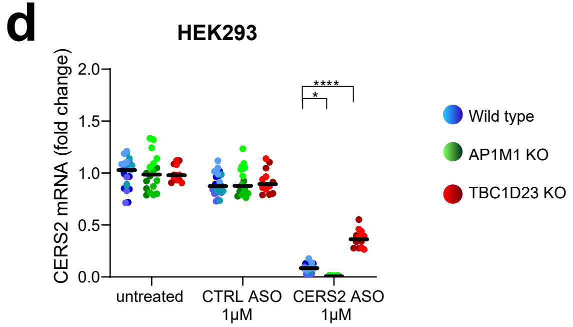


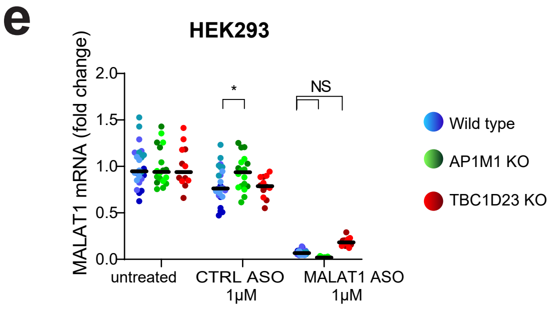
