## Extended data 4 for "A CRISPR/Cas9 screen reveals proteins at the endosome-Golgi interface that modulate cellular ASO activity"

Extended data 4. Malong et al.

**a** U2OS

| Editing frequency of modified CRISPResso2 reads | |
|---|---|
| AP1M1 KO (1) | 99.89% |
| AP1M1 KO (2) | 99.92% |
| TBC1D23 KO (1) | 99.20% |
| TBC1D23 KO (2) | 99.56% |

**b**

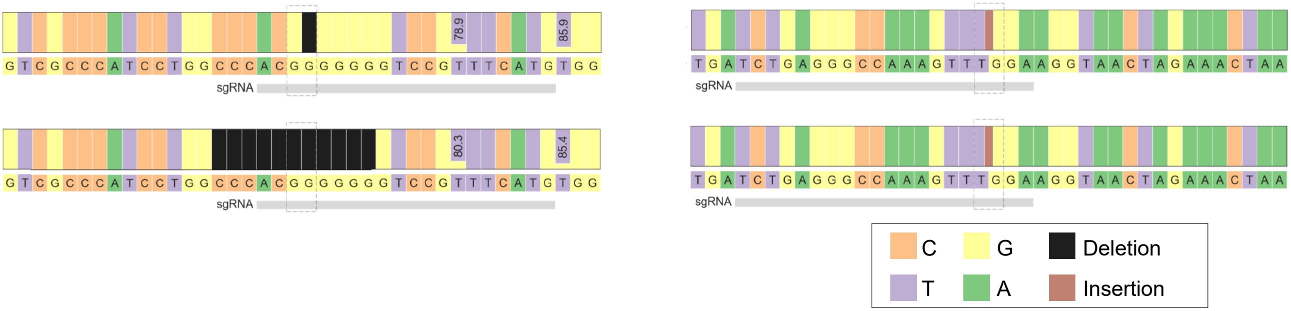


**c**

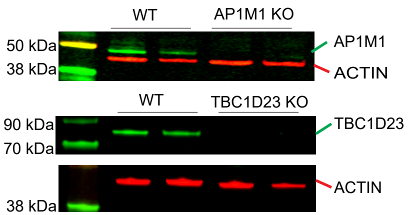


**d**

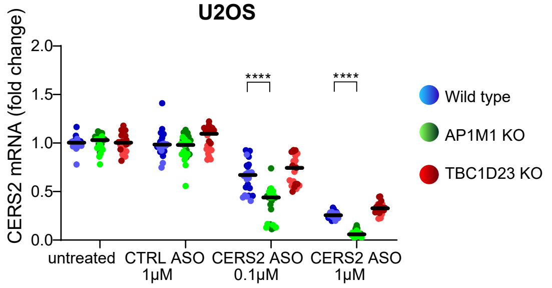


**e**

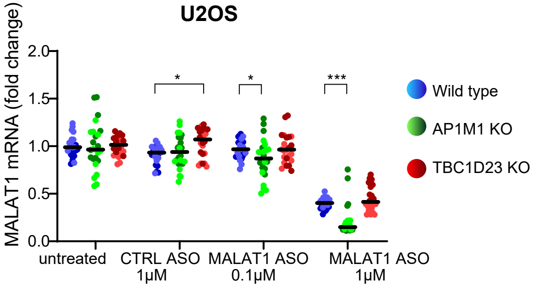
