## Supplementary Tables legends for "A CRISPR/Cas9 screen reveals proteins at the endosome-Golgi interface that modulate cellular ASO activity"

**Malong et al.**

**Supplementary Tables Legends**

**Supplementary Table 1. Nucleotide sequence of the EGFP-splice switch reporter containing mutated intron-2 of human β-globin with additional mutations to reduce background GFP expression.**

Sequence is annotated for GFP, cryptic exon and mutations.

**Supplementary Table 2. SgRNA level analysis of a Genome wide human CRISPR/Cas9 KO screens.**

Table contains the following columns: ID (sgRNA annotation), target (genomic sequence targeted by sgRNA), geneID, geneSymbol. Normalized read counts of sgRNA in all samples (SAM24360646: Reference_R1, SAM24360647: Reference_R2, SAM24360648: Reference_R3 (samples collected before puromycin selection), SAM24360649: Round1_Bulk_R, SAM24360650: Round1_Bulk_R2, SAM24360651: Round1_Bulk_R3 (samples after ASO treatment before FACS sort), SAM24360652: Round1_GFP+_R1, SAM24360653: Round1_GFP+_R2, SAM24360654: Round1_GFP+_R3, SAM24360655: Round1_GFP-_R1, SAM24360656: Round1_GFP-_R2, SAM24360657: Round1_GFP-_R3, SAM24360658: Round2_GFP+_R1, SAM24360659: Round2_GFP+_R2, SAM24360660: Round2_GFP+_R3, SAM24360661:Round2_GFP-_R1, SAM24360662: Round2_GFP-_R2, SAM24360663: Round2_GFP-_R3, with fold change comparisons between populations (LFC_Bulk_to_Reference, LFC_Round2Pos_to_Round2Neg, LFC_Round2Pos_to_Round1Pos, LFC_Round2Neg_to_Round1Neg, LFC_Round1Pos_to_Bulk, LFC_Round1Neg_to_Bulk) with corresponding p-values (Pval_Bulk_to_Reference, Pval_Round1Pos_to_Round1Neg, Pval_Round2Pos_to_Round2Neg, Pval_Round2Pos_to_Round1Pos, Pval_Round2Neg_to_Round1Neg, Pval_Round1Pos_to_Bulk, Pval_Round1Neg_to_Bulk)

**Supplementary Table 3. Gene level analysis of a Genome wide human CRISPR/Cas9 KO screen.**

Table contains the following columns: geneID, geneSymbol; for every comparison between samples Median gene fold change and corresponding rho-value that is a combination of individual sgRNA p-values, p-adjusted value has been calculated.

**Supplementary Table 4.** **Gene ontology analysis for comparison enriched genes of GFP+ Round1 vs unsorted population (Bulk) with a fold change > 1.5 and p-adjusted value < 0.05.**

Table contains the following columns: source (source of the annotation; GO:MF-Gene Ontology Molecular Function branch, GO:BP-Gene Ontology Biological Process branch, GO:CC- Gene Ontology Cellular Component branch, KEGG-Kyoto Encyclopedia of Genes and Genomes pathways, REAC-Reactome pathways); term_name-description of the pathway (group), If not available, repeats the term name; term_id-id in the corresponding source; adjusted_p_value - Hypergeometric p-value after correction for multiple testing; negative_log10_of_adjusted_p_value; term_size - number of genes in the pathway (group); query_size - as ordered query was performed optimal cutoff point for the term was found before the end of the query; intersection_size - The number of genes in the query that are annotated to the corresponding term; effective_domain_size - The total number of genes in the source; intersections - genes in the query that are annotated to the corresponding term.

**Supplementary Table 5.** **Gene ontology analysis for comparison enriched genes of GFP- Round1 vs unsorted population (Bulk) with a fold change > 1.5 and p-adjusted value < 0.05.**

Table contains the following columns: source (source of the annotation; GO:MF-Gene Ontology Molecular Function branch, GO:BP-Gene Ontology Biological Process branch, GO:CC- Gene Ontology Cellular Component branch, KEGG-Kyoto Encyclopedia of Genes and Genomes pathways, REAC-Reactome pathways); term_name-description of the pathway (group), If not available, repeats the term name; term_id-id in the corresponding source; adjusted_p_value - Hypergeometric p-value after correction for multiple testing; negative_log10_of_adjusted_p_value; term_size - number of genes in the pathway (group); query_size - as ordered query was performed optimal cutoff point for the term was found before the end of the query; intersection_size - The number of genes in the query that are annotated to the corresponding term; effective_domain_size - The total number of genes in the source; intersections - genes in the query that are annotated to the corresponding term

**Supplementary Table 6.** **CRISPR/Cas9 knockout arrayed lentiviral screen with high-content imaging readout.**

Table contains the following columns: sgRNA sequences in the corresponding gene, proportion GFP (percentage of GFP+ cells per well) per guide, and per gene (average of all sgRNAs). The not-bold values correspond to the average of the replicates of each sgRNA over two independent experiments. The bold values correspond to the average of the several sgRNA per gene.

**Supplementary Table 7.** **CRISPR/Cas9 knockout arrayed lentiviral screen with FACS readout.**

Table contains the following columns: Gene-sgRNA sequence (gene name and corresponding sgRNA used to target it) and percentage of GFP+ cells (normalized to NT1). The not-bold values correspond to the average of the replicates of each sgRNA over three independent experiments. The bold values correspond to the average of the several sgRNA per gene.

**Supplementary Table 8. Nucleotide sequence of the Piggybac vectors.**

**Supplementary Table 9. Differential expression in HEK293 cells between wild type (WT) and AP1M1 KO. Reported are significant genes with uncorrected p-value < 0.05.**

Table contains the following columns: Contrast (comparison used for analysis), GeneID (EntrezGene), GeneSymbol, logFC (log2 fold change), logCPM (average log2CPM across all samples, CPM stands for counts per million), LR (likelihood ratio of differential expression), PValue (p-value of the generalised linear model fitting with the edgeR method) and FDR (Benjamini-Hochberg adjusted p-values).

**Supplementary Table 10. Differential expression in U2OS cells between wild type (WT) and AP1M1 KO. Reported are significant genes with uncorrected p-value < 0.05.**

Table contains the following columns: Contrast (comparison used for analysis), GeneID (EntrezGene), GeneSymbol, logFC (log2 fold change), logCPM (average log2CPM across all samples, CPM stands for counts per million), LR ( likelihood ratio of differential expression), PValue (p-value of the generalised linear model fitting with the edgeR method) and FDR (Benjamini-Hochberg adjusted p-values).

**Supplementary Table 11. RNA-seq expression matrix (tpm expression values) of the U2OS and HEK293 cells treated with non-targeting control ASOs for three days (1 µM, sequences of CTRL1 and 2  are in methods).**

The table contains the following columns: Ensembl ID, and gene symbol, for every sample. Each condition has 4 replicates.
